# The roles of Transposable Elements and Gene Family Dynamics in Shaping Diversity and Evolution in Diatoms

**DOI:** 10.1101/2025.10.24.684338

**Authors:** Tianze Zheng, Xu Zhang, Tianren Liu, Xinzhu Liu, Chris Bowler, Xin Lin

## Abstract

Diatoms are key players in aquatic ecosystems, having evolved through secondary endosymbiosis. Using long-read sequencing, we investigated how transposable elements (TEs) and gene family dynamics have shaped diatom diversification from inter-lineage to intra-species scales. Across diatom lineages, we identified ecological adaptation-linked expansions, including polyamine synthesis genes for silicification and glutathione S-transferases for oxidative stress resistance. Centric diatoms showed lineage-specific expansion of flotation-associated microtubule genes, while pennate diatoms expanded motility-related actin and myosin genes. At the intra-species level, distinct *Phaeodactylum tricornutum* strains revealed genomic adaptations correlated with their unique features, including strain-specific expansion and contraction in the cruciform strain’s morphological genes and the Baltic Sea isolate’s amine metabolism genes. Our estimates of major lineage divergence times in diatoms (∼202 Ma and ∼173 Ma) were highly consistent with the two deep WGD events (∼200 Ma and ∼170 Ma). At these evolutionary nodes, gene families showed extensive lineage-specific expansions and contractions, likely linking ancient polyploidy to subsequent gene content evolution. Substantial TE expansions occurred more recently (0.5–5 Ma), with most diatoms showing recent bursts of Long Terminal Repeat Retrotransposons (LTR-RTs) and araphid pennate diatoms displaying more ancient TE insertion peaks. This likely reflects the progressive loss of ancient TE copies, leaving only recent TE insertions detectable. Our findings provide genomic evidence for the adaptive evolution of diatoms, highlighting the crucial roles of TEs and gene family dynamics in shaping their morphological diversity and environmental adaptations, and suggesting a potential connection between WGDs, gene family dynamics, and TE insertions in genome evolution.

## INTRODUCTION

Diatoms are a group of diverse single-cell algae general distributed throughout marine and freshwater ecosystems and contributing approximately 20% of the earth’s primary production (Falkowski, Barber, and Smetacek 1998). Diatoms belong to stramenopiles are believed to have undergone secondary endosymbiosis, distinguishing them from higher plants and green algae that evolved from primary endosymbiosis (Falciatore et al. 2020a). Fossil records suggest that diatoms first appeared during the Cretaceous (Bryłka et al. 2024; Roberts, Siepielski, and Alverson 2024). Diatoms are generally classified as centric or pennate (Round, Crawford, and Mann 1990a). Centric diatoms typically have silica frustules that are radially or cylindrically shaped, lacking a raphe, and exhibiting limited mobility (Sims, Mann, and Medlin 2006). Pennate diatoms exhibit bilateral symmetry and typically possess a raphe that enables motility (Cohn and Weitzell 1996). Among pennate diatoms, *Phaeodactylum tricornutum* has emerged as a model species due to its morphological plasticity, physiological adaptability, and availability of multiple natural strains (Bowler et al. 2008a). Previous studies have revealed extensive genetic and transcriptomic diversity among its strains (Rastogi et al. 2020a) (Chaumier et al. 2024).

Comparative genomics provides a powerful framework to investigate genome architecture, genetic diversity, and evolutionary mechanisms across species (Graves 1998). With the advent of long-read sequencing, which produces highly complete genome assemblies, it has become possible to systematically study lineage-specific adaptations and evolutionary mechanisms (Athanasopoulou et al. 2021). Recent advances in plant comparative genomics are exemplified by genomic analyses elucidating the origin of *Citrus* species (Wang et al. 2018), in-depth studies of *Allium fistulosum* (Liao et al. 2022), and the assembly of the *Malus baccata* genome, which revealed adaptive gene family evolution (Chen et al. 2019). Short-read comparative analyses have revealed genomic signatures underlying marine-to-freshwater transitions in diatoms (Roberts et al. 2023). But the contributions of gene family expansions and contractions to diatom diversification remain poorly understood.

In eukaryotes, whole-genome duplication (WGD), is a major mechanism generating new templates for evolutionary innovation (Koszul and Fischer 2009; Lynch and Force 2000) with biased retention of dosage-sensitive genes (e.g., transcription factors) typically causing gene family expansions/contractions (Lockton and Gaut 2005). Transposable elements (TEs) are recognized as major drivers of genome architecture and eukaryotic evolution by introducing mutations and genetic novelty (Gilbert, Peccoud, and Cordaux 2021). However, characterizing and comparing TEs using short-read sequencing remains challenging due to difficulties in resolving repetitive flanking regions essential for accurate TE detection (Anisimova 2019). Long-read sequencing technologies such as PacBio and Oxford Nanopore (ONT) provide critical advantages in resolving repetitive sequences, thereby enabling more comprehensive TE identification and revealing hidden genomic diversity, such as structural variants and TE-rich regions (Warburton and Sebra 2023). Recent studies using long-read sequencing have uncovered extensive TE repertoires and their regulatory roles across eukaryotes (Kapusta and Suh 2017; Stritt et al. 2020; Zhang et al. 2019).

Previous studies, particularly in plants, have shown that TE bursts often follow WGD, a pattern initially framed by McClintock’s “genome shock” hypothesis, which proposed that WGD could trigger immediate TE activation (McClintock 1984). More recent studies, however, suggest that TE accumulation was often a slower, long-term process shaped by relaxed purifying selection in polyploids (Baduel et al. 2019) (Monsen et al. 2025). TE proliferation was also frequently associated with rapid genome size increases, which may occur independently or in synergy with WGD (Chandler and Brendel 2002).

Diatoms have experienced multiple ancient whole-genome duplications (Parks et al. 2018), yet how these events interact with subsequent TE proliferation and gene family evolution remains unclear. Addressing these gaps requires integrating long-read sequencing with comparative genomic approaches across multiple diatom lineages, which offers a unique opportunity to examine how these processes collectively drive ecological diversification in diatoms. Based on the above background, the scientific question of this study is: (1) How do TE landscapes vary across diatom lineages? (2) how do gene family dynamics influence adaptive evolution in diatoms, and what gene families have expanded or contracted in association with ecological adaptations in diatoms? (3) What is the relationship between WGDs, gene family dynamics, and TE proliferation?

In this study, we compiled long-read sequencing data from 29 diatoms and representative outgroups (one coccolithophore, three green algae, and four red algae) to analyze TE diversity and gene family expansion/contraction across lineages. In parallel, we generated ONT-based genomes for three geographically distinct *P. tricornutum* strains to examine how TE dynamics and gene family variation contribute to functional and physiological differentiation within a model species. We further investigated the relationships between WGDs, gene family dynamics, and TE proliferation to understand their coordinated roles in shaping genome evolution. Together, this comparative genomics with strain-level analyses provides new insights into the evolutionary forces shaping diatom diversity.

## MATERIALS AND METHODS

### Genome sequencing of *P. tricornutum*

Three *P. tricornutum* strains—PtSCS, PtECS, and Pt4—were sequenced (**Fig. 1**). Pt4 was isolated from the Baltic Sea and deposited in the Culture Collection of Algae and Protozoa (CCAP 1052/6) (Rastogi et al. 2020a). PtSCS was collected from the South China Sea in 2004 and deposited in the Center for Collections of Marine Bacteria and Phytoplankton, Xiamen University (CCMBP106) (Huang et al. 2020). PtECS, a cruciform strain from the Yangtze River estuary in the East China Sea, was deposited in the same collection (CCMBP267).

**Fig. 1.**
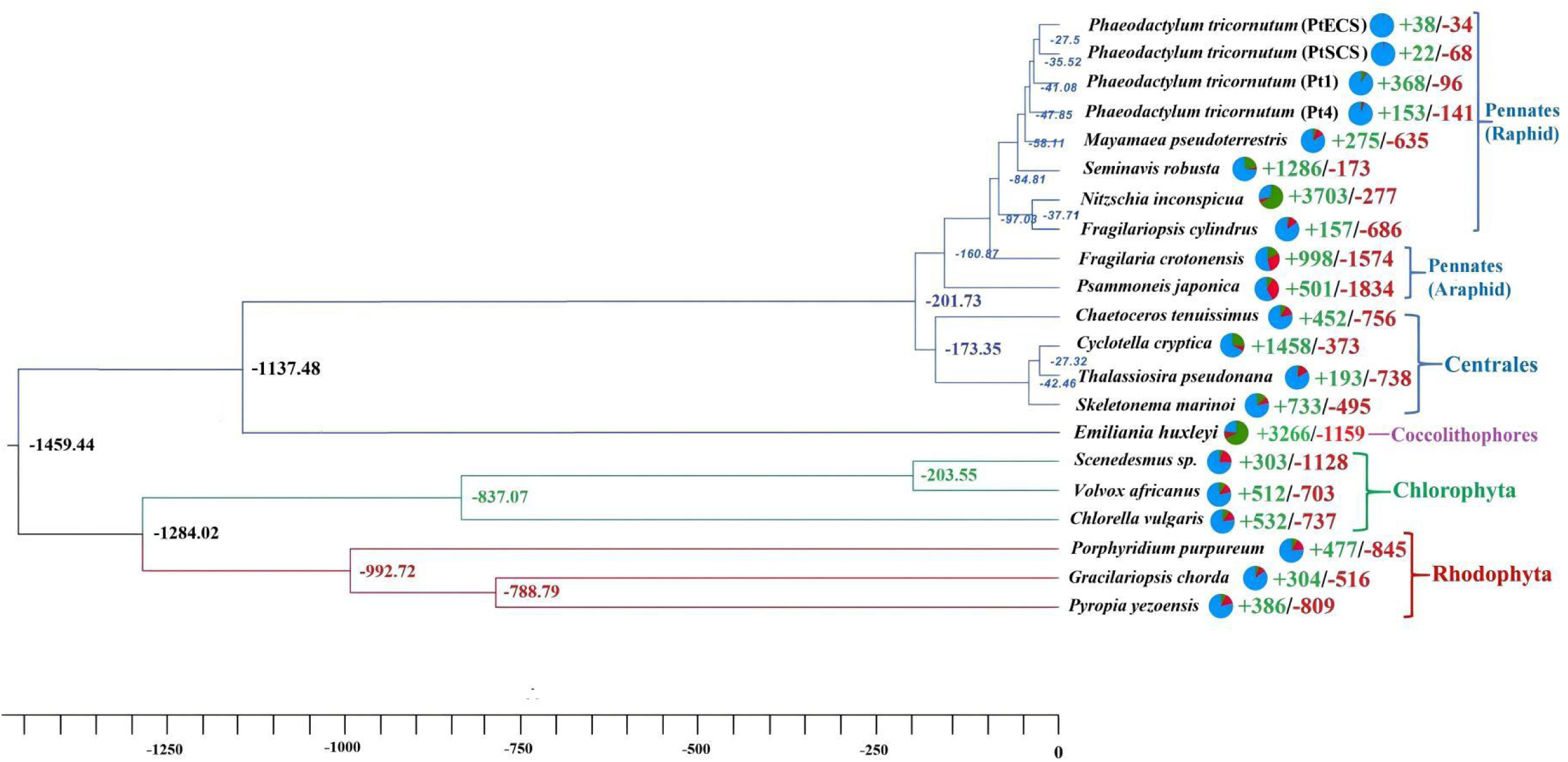
Analysis of divergence times and gene family evolution in 14 diatom species, based on long-read sequenced, well-annotated nuclear genomes from both pennate and centric lineages, one coccolithophore, three green algae, and three red algae. Different phyla are color-coded. The expansion (+) and contraction (-) of gene families are displayed on the phylogenetic branches. Pie charts following species names show contraction (red), expansion (green), and unchanged (blue) gene families, while numbers following each node indicate estimated divergence times for those taxa. One unit of length equals one million years.

Cultures were treated with an antibiotic cocktail, maintained in f/2 enriched seawater at 19 °C under a 12 h light/dark cycle, and harvested during exponential growth. High-quality genomic DNA was extracted following (Filloramo et al. 2021): cells were harvested by centrifugation, lysed in SDS buffer, subjected to ten freeze–thaw cycles, digested with proteinase K, treated with RNase A, purified by phenol–chloroform extraction and isopropanol precipitation, and finally resuspended in 10 mM Tris-HCl (pH 8.0).

Sequencing was performed at Biomarker Technologies (Beijing, China) using Oxford Nanopore Technology (ONT). Long reads (up to ∼2 Mbp) were generated with the SQK-LSK109 ligation kit. Raw signals were collected using MinKNOW and subsequently base-called with Guppy (Oxford Nanopore Technologies; specific version information not available from archived data), and converted to FASTQ format for downstream analysis (Deamer, Akeson, and Branton 2016; Warburton and Sebra 2023).

### Genome assembly and annotation

We conducted *de novo* genome assembly using filtered reads, employing NextDenovo followed by NextPolish-based long-read correction (two rounds) and Pilon-mediated short-read polishing (three rounds) with Illumina data. Gene structural annotation was conducted using Braker2 (Brůna et al. 2021) and EVidenceModeler (EVM) processes, to incorporate transcriptomic data. Additionally, gene function annotation was conducted using EggNOG-mapper (Huerta-Cepas et al. 2017).

### Genomic data compiled and analysis of long-read sequencing data

We focus primarily on diatom species with available high-quality long-read genome assemblies. Red algae (Rhodophyta) and green algae (Chlorophyta) were included as outgroups for comparative purposes, serving to highlight lineage-specific features in diatoms and distinguish them from ancestral or broadly conserved genomic traits. The use of outgroups allows for a more nuanced interpretation of diatom evolution in the broader context of plastid-bearing algae. We analyzed published genomic data from 29 diatoms sequenced with Nanopore/PacBio platforms, including the *P. tricornutum* strain Pt1 isolated from the British Irish Sea (Filloramo et al. 2021). The dataset also incorporated one coccolithophore, three green algae, and four red algae obtained from NCBI, JGI, and Ensembl databases. We performed comparative genomic analysis on selected diatoms and outgroups (one coccolithophore, three green algae, and three red algae) with high-quality genomes and GFF annotations (**Table 1**). Genome completeness was assessed using BUSCO v5.4+ with both universal eukaryotic (eukaryota_odb10) and lineage-specific datasets: stramenopiles_odb10 for diatoms, chlorophyta_odb10 for green algae, and rhodophyta_odb12 for red algae, ensuring the most appropriate evolutionary context for each taxonomic group. BUSCO completeness assessment revealed generally high genome quality across the analyzed species, with scores ranging from 76.0% to 100%. Most diatom genomes showed excellent assembly quality (92-99% BUSCO), with *Nitzschia inconspicua* being complete at 100%.

**Table 1.**
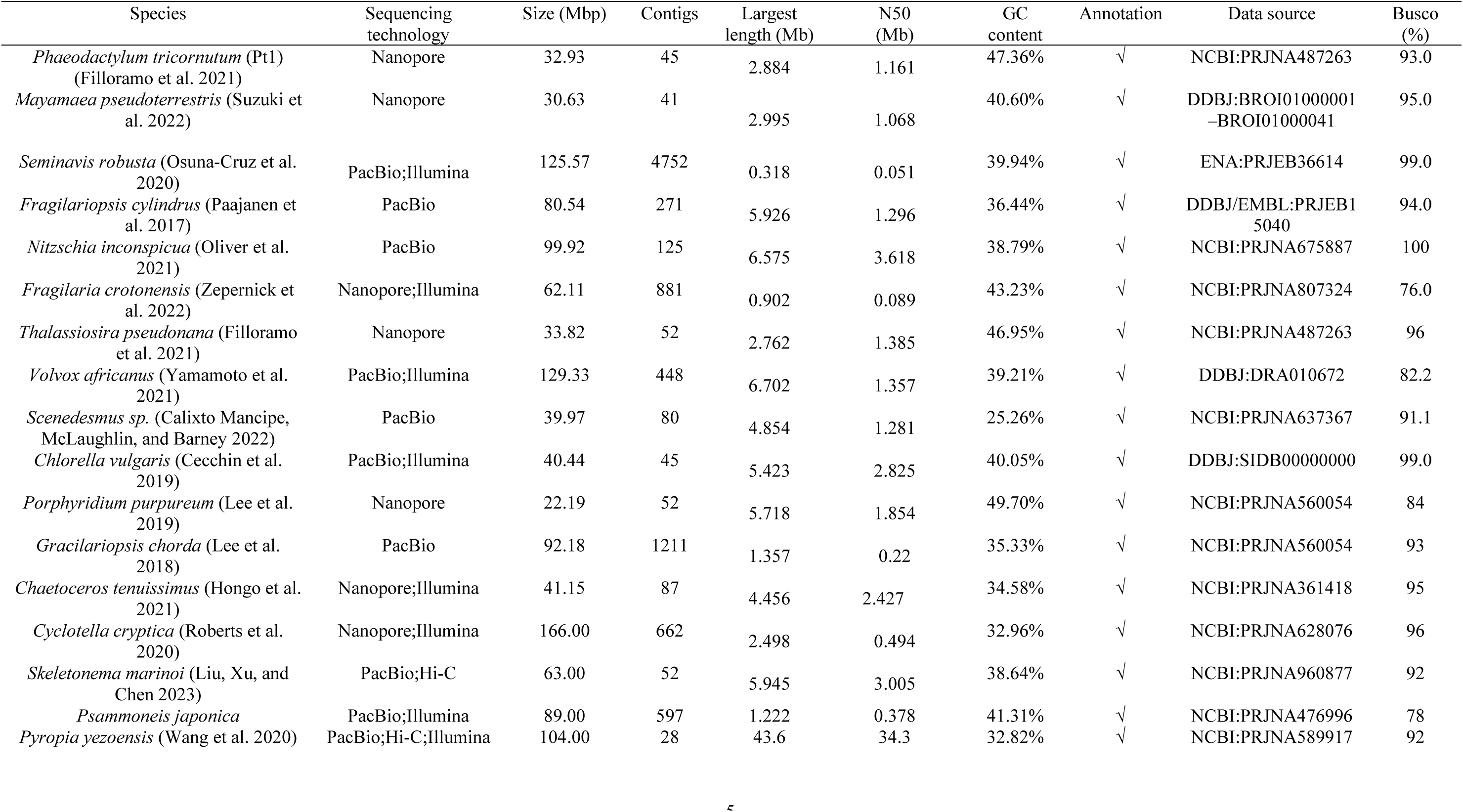
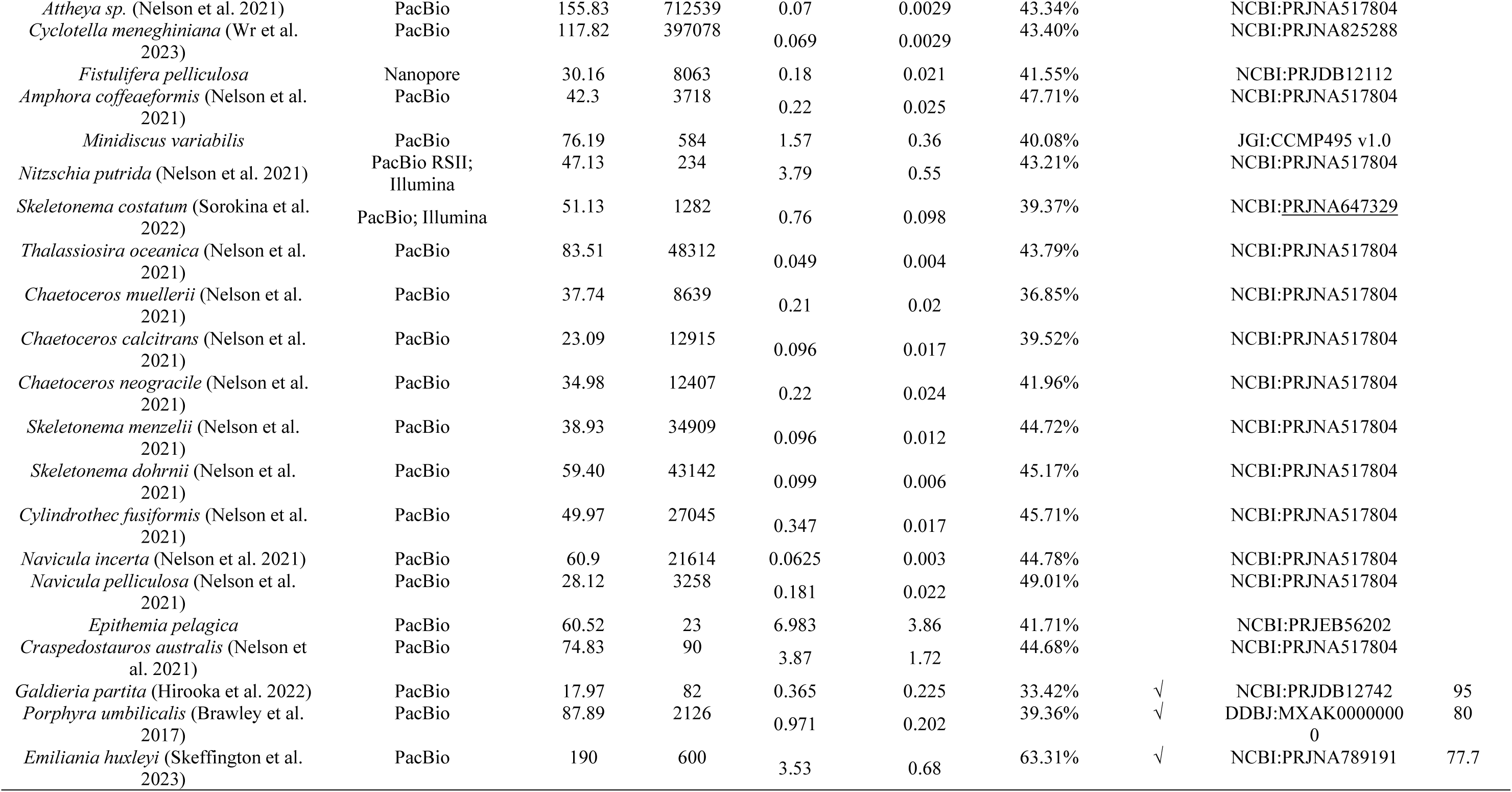
Overview of genomic data from published studies for selected diatoms, red algae, and green algae, including species name, sequencing technology used (e.g., Nanopore, PacBio, Illumina), genome size in megabase pairs (Mbp), number of contigs, the length of the largest contig in Mbp, the sequence length of the shortest contig at 50% of the total assembly length (N50) in Mbp, GC content percentage, Data source, and the score of Busco.

### Comparative genomic analysis of diatoms and selected coccolithophore, red and green algal species

We conducted a comparative genomic analysis of 21 species, including 14 diatoms with high-quality annotated genomes (pennate and centric), one coccolithophore, three green algae, and three red algae as outgroups. The diatoms included four *P. tricornutum* strains (Pt1, Pt4, PtSCS, PtECS). We performed gene family clustering analysis using Diamond to compare sequences and identify homologous proteins, followed by OrthoFinder (Emms and Kelly 2019) to cluster gene families and identify shared and unique ones.

We reconstructed the phylogeny and estimated divergence times using a Bayesian approach in MCMCtree (PAML package; Dos Reis 2022). The analysis was based on OrthoFinder-derived alignments filtered with TrimAl (gt=0.6, cons=60), with the species tree constructed using RAxML-HPC-PTHREADS and rooted in MEGA. Divergence times were estimated using the MCMCtree tool. Gene family expansions and contractions were analyzed using CAFE5 (Mendes et al. 2021).

We performed Gene Ontology (GO) and Kyoto Encyclopedia of Genes and Genomes (KEGG) enrichment analyses on expanded/contracted gene families across different evolutionary nodes. In addition to analyses at key evolutionary nodes, we also conducted species-specific GO and KEGG analyses for the model diatom *P. tricornutum*, *N. inconspicua* (selected due to its extensive gene family expansion), and the two araphid pennate diatoms. The araphid pennate diatom *Psammoneis japonica* and *Fragilaria crotonensis* displayed unique genomic and TE features, suggesting their potential transitional position in diatom evolution between centric and pennate diatoms. To explore their evolutionary relationship within the centric-pennate diatom divergence, we performed whole-genome duplication (WGD) analysis using the ‘WGD’ tool, calculating synonymous substitutions per synonymous site (Ks) (Zwaenepoel and Van de Peer 2019).

### TE annotation and divergence analysis of diatoms and representative red algal and green algal species

TE annotation was performed using a sequential, integrated pipeline. First, comprehensive *de novo* TE identification and initial classification were conducted using the Extensive *de novo* TE Annotator (EDTA) (Su et al. 2021), which detects TEs based on structural features including terminal repeats, target site duplications, and homology. Second, DeepTE (Yan, Bombarely, and Li 2020), a deep learning-based classifier, was applied to refine the classification of TEs that remained unclassified or ambiguously annotated by EDTA. This complementary approach maximizes both the sensitivity of TE detection and the accuracy of superfamily assignment. Based on the integrated annotation results, we compiled characteristic parameters of TEs across different species of diatoms, green algae, and red algae.

We selected twelve diatom species for a comparative study of TE evolutionary dynamics. The selection encompassed major phylogenetic lineages and was based on high-quality genome assemblies, diverse TE content, key evolutionary positions, and distinct gene family dynamics. We dated Long Terminal Repeat Retrotransposons (LTR-RTs) insertion time using a sequence identity-based consensus approach, where the nucleotide identity between individual TE copies and their subfamily consensus sequences serves as a proxy for time since insertion. To further place these estimates on an absolute timescale, we applied a molecular clock calibration based on the mutation rate of *P. tricornutum* (Krasovec, Sanchez-Brosseau, and Piganeau 2019). Using RAxML, we constructed a phylogenetic tree based on the RT domains of full-length LTR-RTs.

## RESULTS

### Estimation of divergence time and gene family expansions and contractions among diatoms

To reconstruct diatom evolutionary history, we analyzed 21 genomes, including 14 diatoms representing major centric and pennate lineages, alongside one coccolithophore, three green algae, and three red algae. Divergence time estimates place the origin of diatoms at ∼202 Mya, with centric diatoms emerging earlier (∼173 Mya) and pennate diatoms diversifying later, including *Psammoneis japonica* (∼161 Mya) and *Fragilaria crotonensis* (∼97 Mya), consistent with previous evolutionary frameworks (Falciatore et al. 2020b).

In diatoms, glutathione metabolism genes—particularly glutathione S-transferases—showed widespread expansion, as did polyamine metabolism and carotenoid biosynthesis pathways, both central to frustule silicification and photoprotection. In centric diatoms, gene families related to microtubules and cellular protrusions were expanded. Pennate diatoms displayed expansions of actin- and myosin-related genes underlying motility (**Supplementary Table S2**).

The pennate araphid diatoms *P. japonica* and *F. crotonensis* showed substantial gene family contractions (1,834 and 1,574, respectively). *C. cryptica* expanded 1,458 gene families, while *S. robusta* and *N. inconspicua* expanded 1,286 and 3,703 gene families, respectively (**Fig. 1**). Functional enrichment pointed to expansions of fucoxanthin chlorophyll a/c binding proteins, HSP70s, carbonic anhydrases, and Rubisco at key evolutionary nodes (**Supplementary Table S5**). We observed a striking gene family expansion in *N. inconspicua*, with functional enrichment analyses revealing that the expanded families were primarily associated with spliceosome, ribosome, RNA metabolism, and cell cycle regulation (KEGG), as well as nitrogen compound transport, protein localization, energy metabolism, and organelle organization (GO) (**Fig S3, S4**). In *P. tricornutum*, genes involved in DNA integration and histone modifications showed pronounced expansions, suggesting lineage-specific regulatory innovations (**Supplementary Table S2**).

### The diversity of TEs in diatoms and comparison with other algal species

To examine TE distribution patterns across algal lineages, we analyzed and compared TE content in diverse diatoms, green algae, and red algae. *Cyclotella cryptica* had the highest overall TE and LTR-RT proportions among diatom genomes (53.37% and 34.98%, respectively). *Attheya* sp. ranked second in total TE content (47.84%) and displayed the highest proportion of TIR elements (15.84%) among all species analyzed (**Fig. 2**; **Supplementary Table S1**). *Chaetoceros australis* had the second-highest LTR-RT proportion (17.95%).

**Fig. 2.**
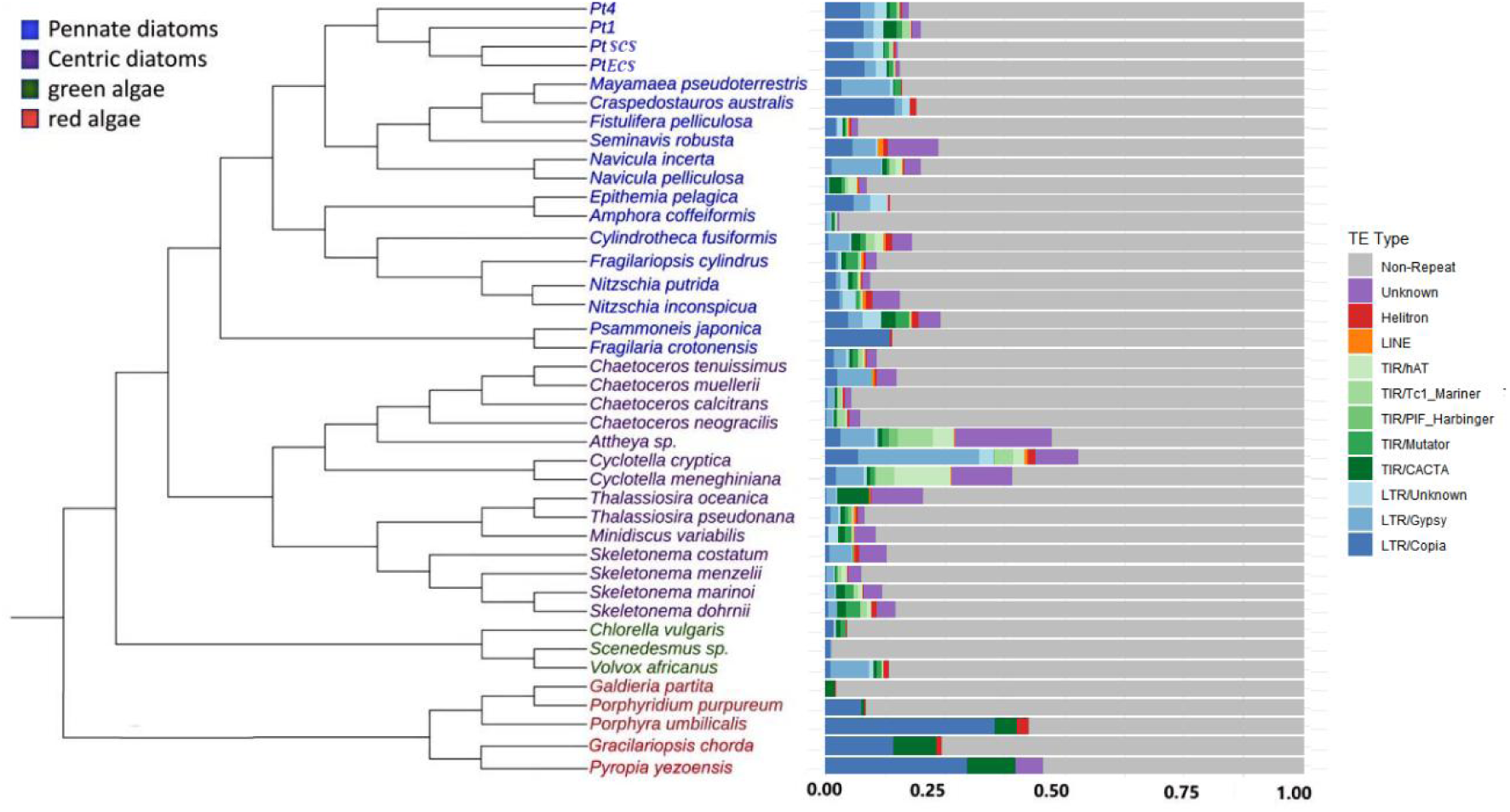
The content of TEs in 32 species of diatoms, three green algae and five red algae species. The left side of the figure shows the phylogenetic relationship of centric diatoms, pennate diatoms, green algae, and red algae, represented in different color areas; the right side represents the proportion of TEs in the genome of each species, with different colors distinguishing different types of TEs (subdivided into superfamilies). The phylogenetic relationship between different diatoms and other algae species is based on NCBI common tree (https://www.ncbi.nlm.nih.gov/Taxonomy/CommonTree/wwwcmt.cgi).

Species with larger genomes, such as *C. cryptica* (166 Mb) and *Attheya* sp. (155.8 Mb), contained substantially higher proportions of TEs compared to species with smaller genomes. Notably, *F. crotonensis*, an araphid pennate diatom, exhibited distinctive TE distribution patterns with LTR-RTs accounting for up to 90% of its total TE content.

### Phylogenetic analysis of LTR-RTs indicates divergent insertion patterns among diatoms

We conducted a phylogenetic analysis using conserved domain sequences to elucidate the distribution and evolution of LTR-RTs across different diatom species. *C. cryptica*, *F. crotonensis* and *P. japonica* exhibited high LTR-RT diversity. In contrast, *Thalassiosira pseudonana* exhibited markedly lower LTR-RT diversity (**Fig. 3**).

**Fig. 3.**
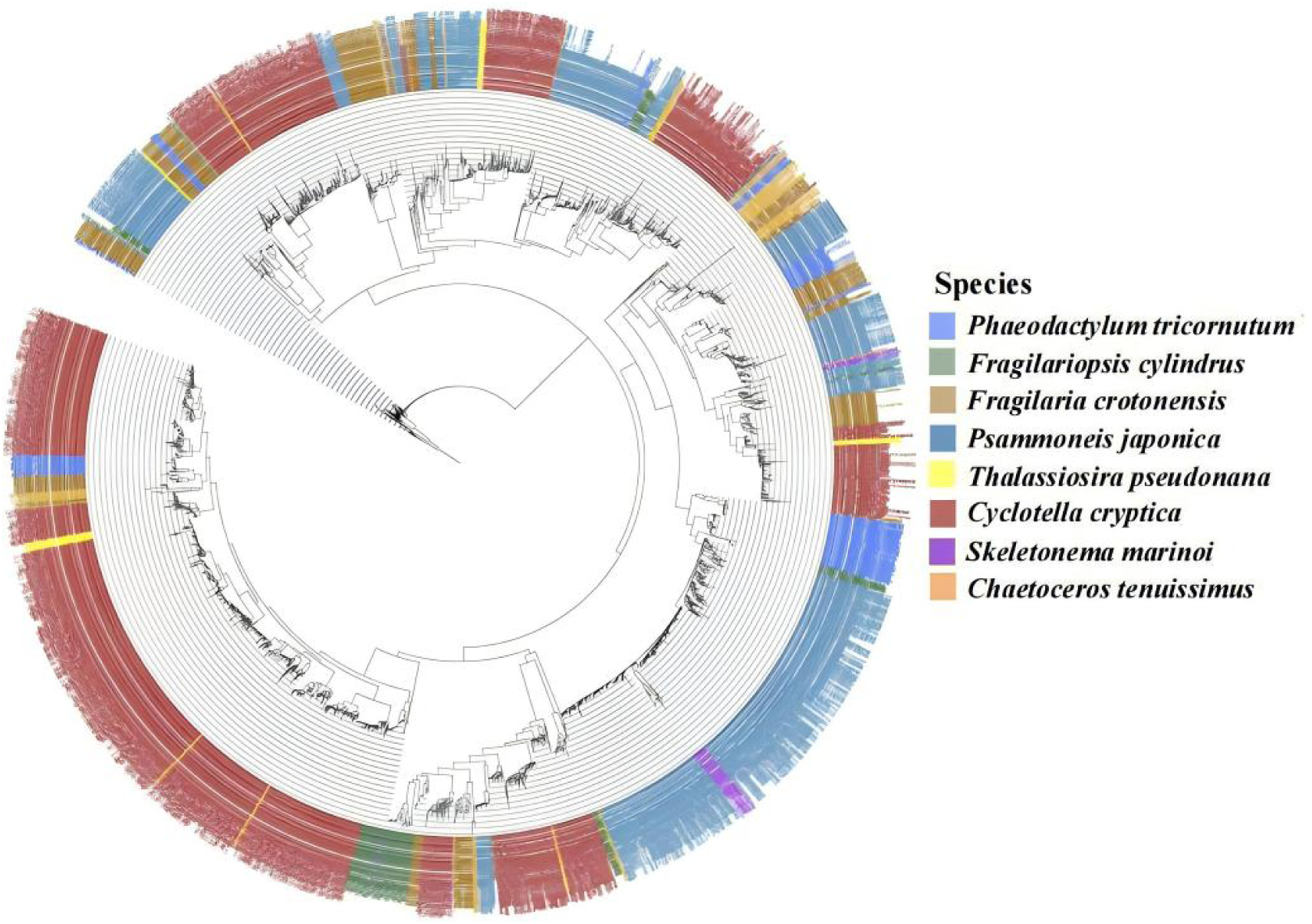
The phylogenetic analysis of LTR-RTs across eight diatom species (including five representative pennate and centric species) based on conserved domain sequences. The tree was constructed using conserved RT domain sequences. Different colored labels represent different diatom species.

### Diversity in LTR-RT insertion patterns among diatoms

To investigate TE dynamics in diatoms, we analyzed LTR-RT insertion patterns in species with high TE content and high-quality genome assemblies. Insertion times were estimated using sequence identity between LTR-RT elements and their ancestral sequences based on EDTA annotations. Higher sequence identity indicates more recent insertions, whereas lower identity reflects older events that have accumulated mutations over time.

Most diatoms exhibited a recent burst of LTR-RT activity. *Attheya* sp. showed the most prominent recent peak, while *C. cryptica* and *Psammoneis japonica*—despite their high TE content—lacked detectable recent insertions. Interestingly, the araphid diatoms *F. crotonensis* and *P. japonica* displayed insertion patterns consistent with more ancient evolutionary events (**Fig. 4**).

**Fig. 4.**
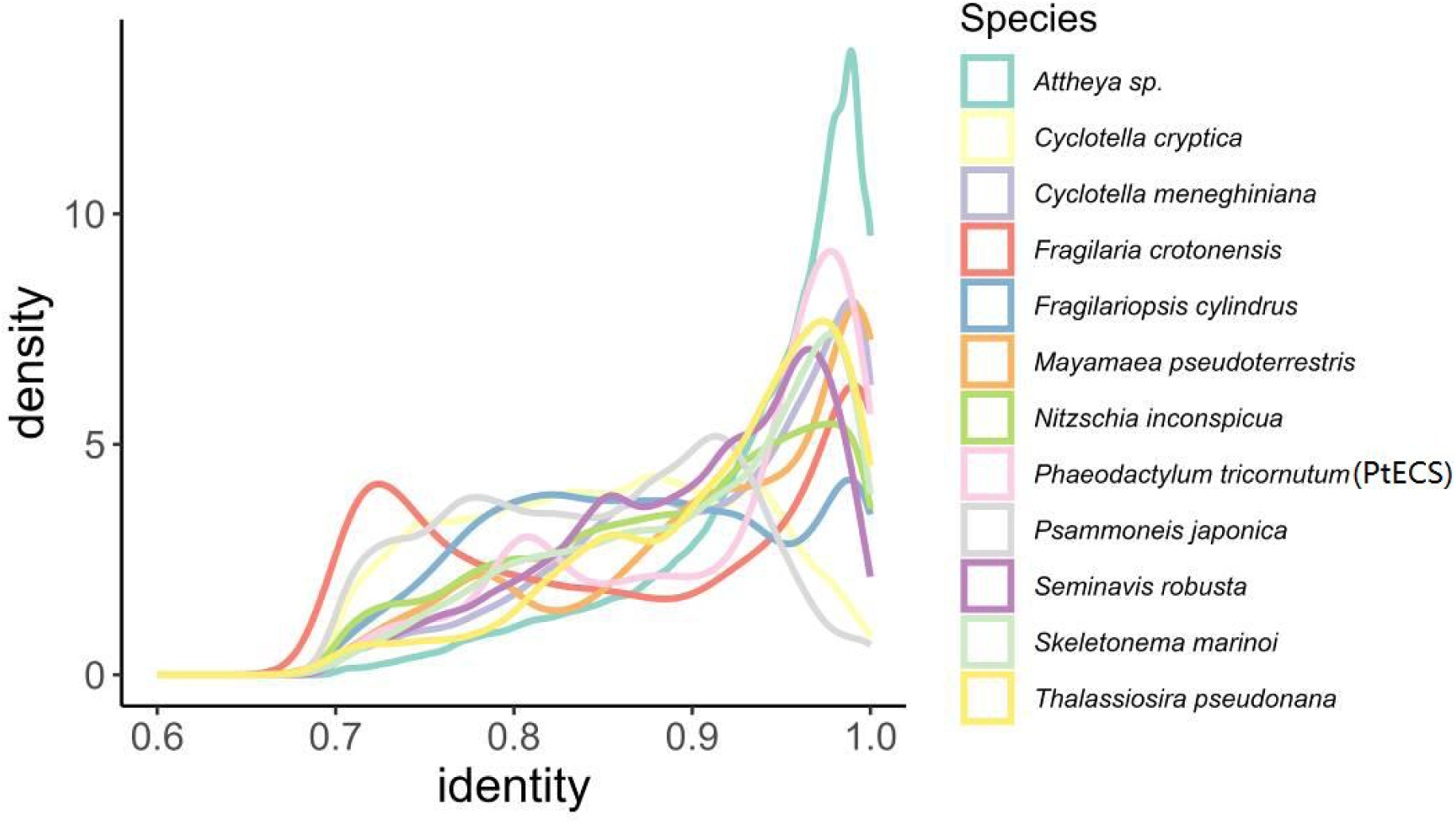
Density plot of sequence identity between LTR-RT of diatoms. The X-axis shows identity, reflecting how genetically similar the LTR sequences are to ancestral ones. A higher identity score signifies closer resemblance to the ancestral sequences, suggesting a more recent divergence. The Y-axis depicts the density, representing the frequency of LTR-RT elements sharing an identity level (indicated by the X-axis value). Peaks on the graph signify widespread transposon replication events across the genome.

Based on the mutation rate of *P. tricornutum* (Krasovec et al. 2019), we inferred the insertion time distributions of Copia and Gypsy LTR-RTs across six diatom genomes (**Fig. 5**). The results revealed pronounced recent expansions with distinct patterns among species: in *C. cryptica* and *T. pseudonana*, Gypsy elements dominated with expansions extending to ∼3–4 Mya; *F. crotonensis* and *P. japonica* showed continuous Gypsy accumulation accompanied by localized Copia peaks; while in *N. inconspicua* and *P. tricornutum*, expansions were almost exclusively driven by Copia with more recent insertions (∼0.5–1.5 Mya).

**Fig. 5.**
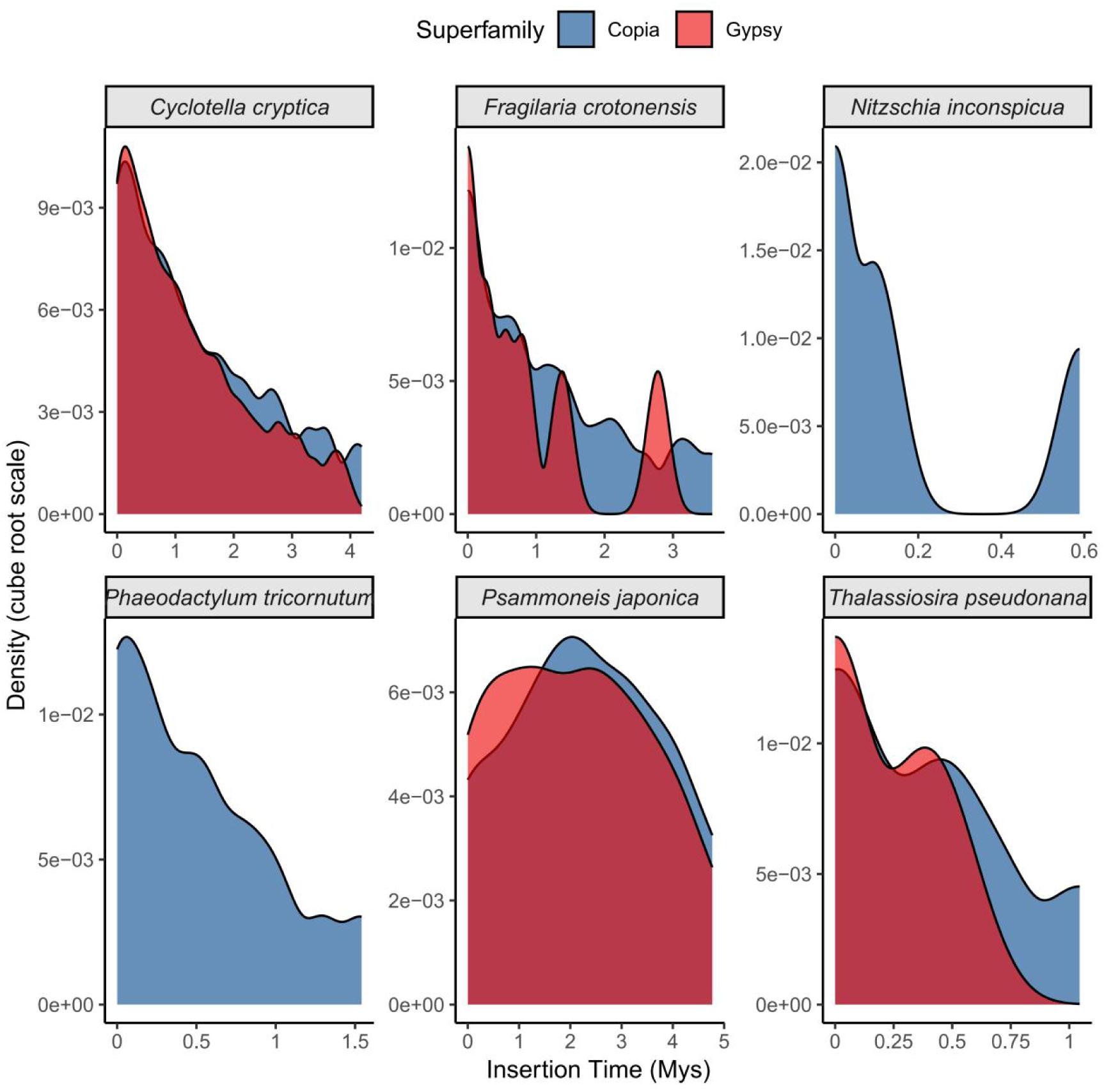
Insertion time distributions of Copia and Gypsy retrotransposons across six diatom genomes based on the mutation rate of *P. tricornutum* (Krasovec et al. 2019)

### Genome comparison of different *P*. *tricornutum* strains

To compare genomic features across *P. tricornutum* strains, we analyzed their genome assemblies and structural variations. Relative to the Pt1 reference genome (Bowler et al. 2008b), Pt4 harbored the highest number of structural variations (5,520), followed by PtSCS (4,974) and PtECS (3,647). Assembly statistics also differed: Pt4 contained the largest number of contigs (68), whereas PtSCS and PtECS had 30 and 36, respectively. The longest contigs were slightly larger in PtSCS and PtECS (2.81 Mbp) than in Pt4 (2.69 Mbp). PtSCS showed the highest N50 (1.52 Mbp), while PtECS had the lowest (1.3 Mbp). In contrast, GC content was nearly identical across strains (∼48.5%), and TE content was broadly similar, though PtSCS exhibited the lowest proportion (15.6%) (**Table 2**).

**Table 2.** Genome assembly parameters of different *P. tricornutum* strains include average sequencing depth, genome size in megabase pairs (Mbp), number of contigs, the length of the largest contig in Mbp, the sequence length of the shortest contig at 50% of the total assembly length (N50) in Mbp, GC content percentage, TE content, and detected SVs).

| strain | Pt4 | PtSCS | PtECS |
| --- | --- | --- | --- |
| Average depth | 139X | 168X | 99X |
| Data size (Mbp) | 4856.75 | 5362.84 | 3150.76 |
| Genome size (Mbp) | 30.5 | 29.15 | 29.07 |
| Contigs | 68 | 30 | 36 |
| Largest contig length (Mbp) | 2.69 | 2.81 | 2.81 |
| N50 (Mbp) | 1.31 | 1.52 | 1.3 |
| GC content | 48.53% | 48.59% | 48.51% |
| TE content | 17.72% | 15.60% | 16.11% |
| structural variations (SV) | 5520 | 4974 | 3647 |

### Gene family contraction and expansion in different *P. tricornutum* strains

To investigate gene family dynamics across *P. tricornutum* strains, we conducted a comparative genomics analysis and reconstructed their phylogeny using divergence time estimation. Pt4 diverged earliest (∼41.1 Mya), followed by Pt1 (∼35.5 Mya), while PtSCS and PtECS separated more recently (∼27.5 Mya) (**Fig. 6**).

**Fig. 6.**
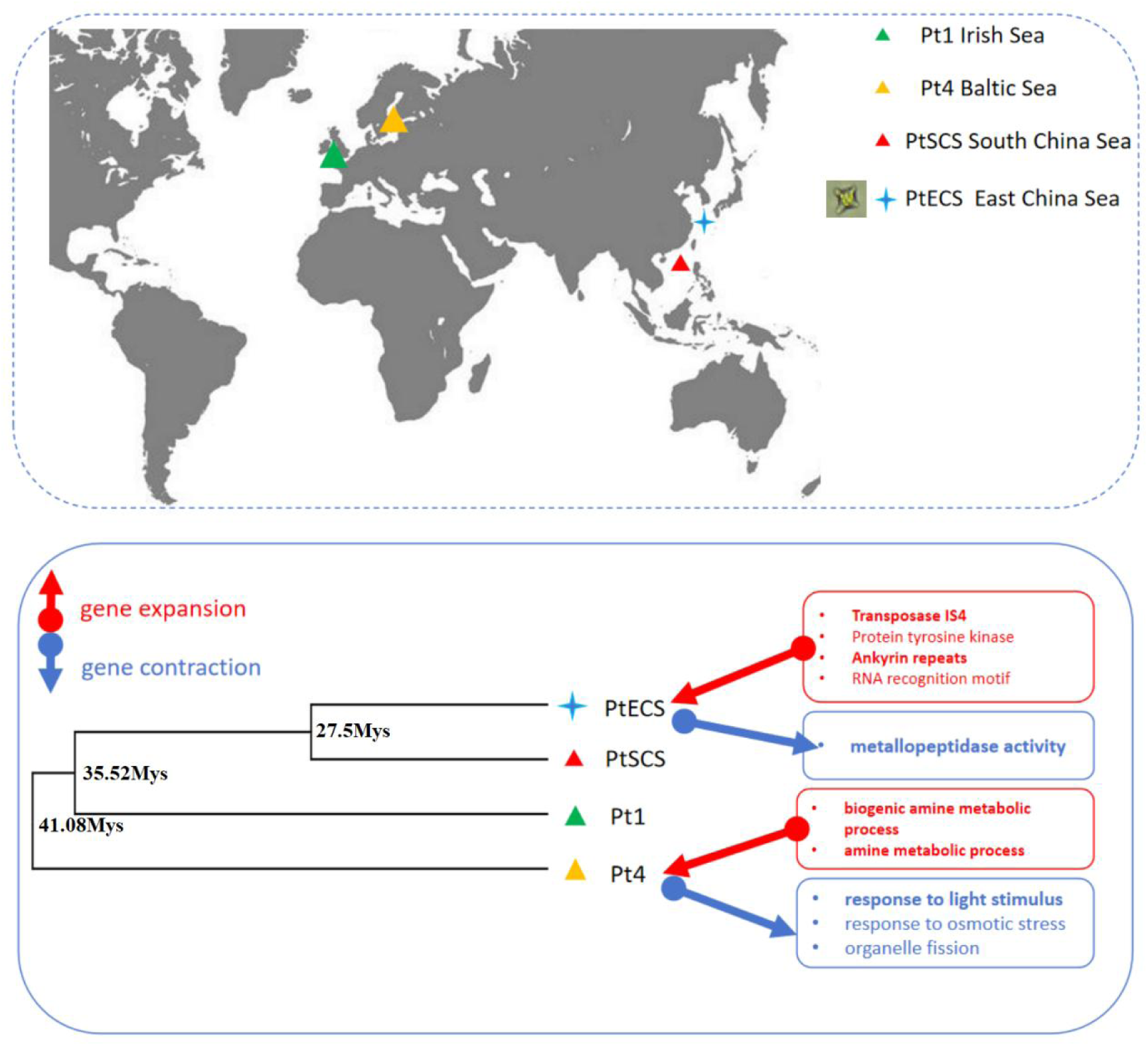
Gene contraction and expansion in different strains *P. tricornutum* isolated from different regions. The upper panel illustrates the distribution of various *P. tricornutum* strains from different locations on the map. Gene functions that have undergone expansion or contraction are indicated in the lower panel with red arrows for expansion and blue arrows for contraction.

Strain-specific patterns of gene family change were evident. In PtECS, transposase IS4, ankyrin repeats, and protein tyrosine kinase families expanded, whereas metallopeptidases contracted. In Pt4, gene families related to amine metabolism expanded, while light response genes contracted (**Fig. 6**). Notably, the pyruvate orthophosphate dikinase (PPDK) family, linked to C_4_ metabolism, expanded in Pt4, whereas other strains showed expansions of genes associated with C_3_ metabolism (**Supplementary Table S4**).

### TE diversity among different *P. tricornutum* strains

We analyzed TE composition across *P. tricornutum* strains using EDTA and DeepTE. LTR-RTs were the most abundant elements, accounting for 12-12.9% of the genome, representing approximately 75–80% of the total TE content, with PtECS having the highest proportion and PtSCS the lowest. Given their genomic dominance and the documented activity of LTR/Copia elements in *P. tricornutum* (Maumus et al. 2009), we focused our subsequent polymorphism analyses primarily on LTR-TEs. TIR elements contributed 1.97–3.21% of the genome, representing the second most abundant TE class, and were therefore also included in our TE insertion analysis (**Fig. 7C**). In contrast, LINEs were rare (<0.5%) and other TE types showed relatively low abundance (**Fig. 7A**). Overall, total TE content remained relatively stable among strains (15.6–17.7%), but our estimates were markedly higher than the 6.4% previously reported for the reference genome (**Fig. 7A**).

**Fig. 7.**
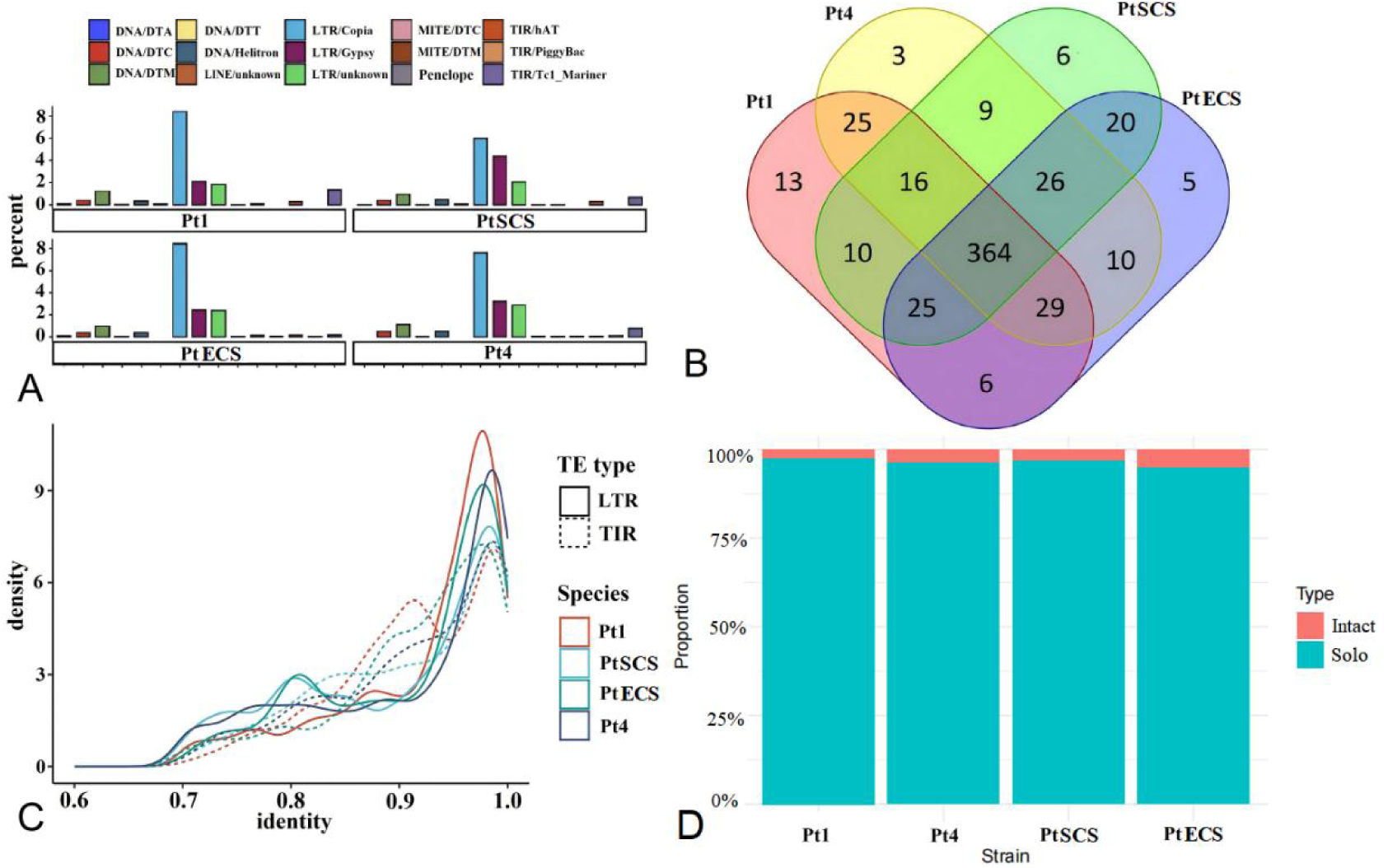
TE diversity and sequence identity across *P. tricornutum* strains. TE proportion of *P. tricornutum* bar graph illustrates the distribution percentages of various transposon types within *P. tricornutum*, with each color signifying a distinct type of TEs (A). LTR/Copia element variability in different strains chart enumerates both the common and unique LTR/Copia elements found in different *P. tricornutum* strains (B). Density plot of sequence identity delineates the sequence identity between transposons from four *P. tricornutum* strains compared to their original sequences (C).

A total of 364 LTR/Copia sequences were shared across all four strains, while strain-specific sequences numbered 13 in Pt1, 3 in Pt4, 6 in PtSCS, and 5 in PtECS. Pairwise comparisons revealed stronger similarity between Pt1 and Pt4 (25 shared sequences) and between PtSCS and PtECS (20 shared sequences) (**Fig. 7B**). All strains exhibited genomic signatures of recent TE activity, but their patterns differed. PtSCS and PtECS showed distinct LTR-RT insertion peaks at 80% sequence identity, whereas Pt1 displayed a pronounced TIR expansion at 90% identity, suggesting a recent large-scale insertion event (**Fig. 7C**). Phylogenetic analysis of *P. tricornutum* LTR/Copia elements (**Fig. 8**) confirmed the seven CoDi groups (CoDi1–CoDi7) described by (Maumus et al. 2009) and identified 30 unclassified elements that likely represent novel retrotransposon families. These novel elements showed strain-specific distributions: 5 in Pt1, 7 in Pt4, and 9 each in the Chinese strains PtSCS and PtECS.

**Fig. 8.**
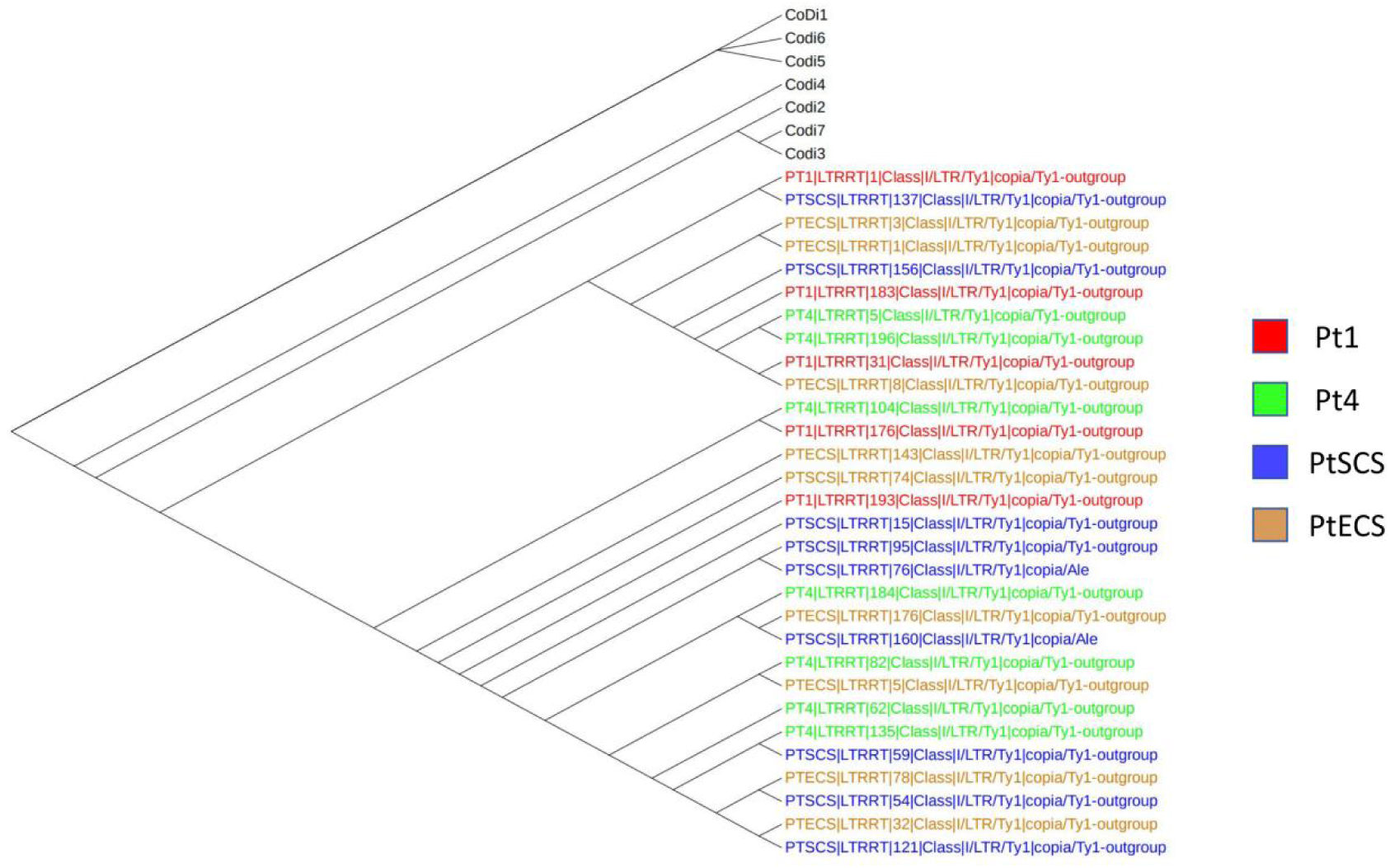
Phylogenetic tree of RT domains from LTR/Copia transposons in *P. tricornutum*, including known typical CoDi groups (CoDi1-CoDi7) and newly discovered LTR/Copia sequences from four strains. Colors represent different strains: red (Pt1), green (Pt4), blue (PtSCS), and brown (PtECS). The phylogenetic tree illustrates the evolutionary relationships between these newly discovered LTR/Copia sequences and known CoDi groups.

Among all strains, the intact LTR-RTs (the incomplete form) constituted a minor fraction of the total LTR-RT content, with the remainder being predominantly solo LTRs (the complete form). The Intact LTR content of the PtECS strain was the highest, and it was also the strain with the latest stage of differentiation (**Fig. 7D**).

## DISCUSSION

### Diatom evolutionary events and the potential association between contraction/expansion of gene families and morpho-physiological traits

Diatoms and coccolithophores have evolved specialized adaptations through gene family expansions that underpin their ecological success (**Fig. 9**). Expansions of fucoxanthin– chlorophyll protein (FCP) genes enhance light harvesting in the blue–green spectrum (450– 570 nm) characteristic of marine environments (Cao et al. 2023). Duplications of carbonic anhydrase and RubisCO genes support CO₂-concentrating mechanisms (CCMs), facilitating efficient carbon fixation under low-CO₂ conditions (Reinfelder 2011). Expansions of heat shock protein (HSP) families further contribute to resilience under thermal stress, a key trait in dynamic marine habitats (Rousch, Bingham, and Sommerfeld 2004).

**Fig. 9.**
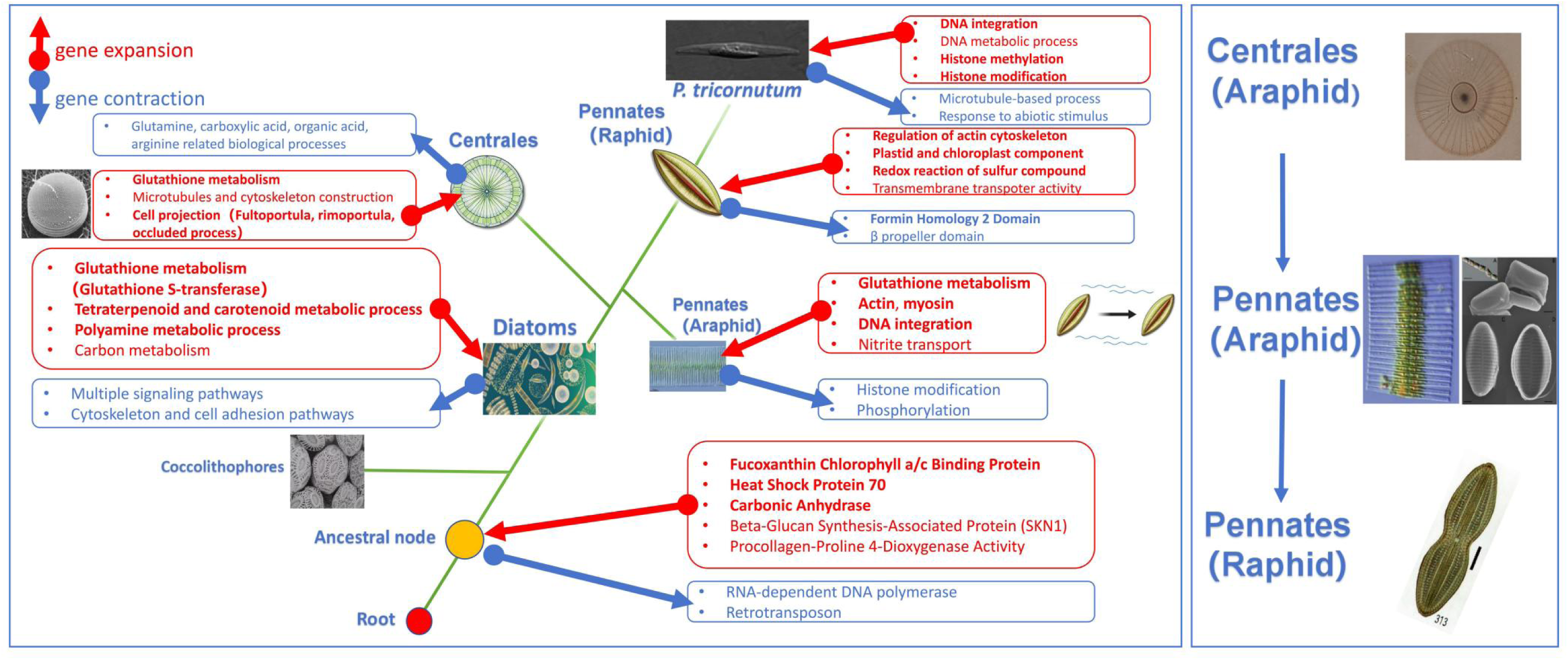
Schematic of gene function contraction and expansion in diatoms based on the whole-genome comparative analysis. In the left panel, red arrows symbolize genes that have expanded, while blue arrows represent genes that have contracted. Green lines denote evolutionary relationships. The descriptions inside the boxes provide a general summary of the gene functions that have contracted or expanded at the current node compared to its ancestral node. Some image materials are from Flickr (https://www.flickr.com/photos/tags/flickr/)

Diatoms exhibited notable gene family expansions in polyamine metabolism, essential for silica frustule formation—a defining adaptive trait with major ecological significance (Poulsen, Sumper, and Kröger 2003). They also expanded glutathione metabolism genes, particularly glutathione S-transferases (GSTs), which are central to oxidative stress defense, detoxification, and cellular signaling (Aloke, Onisuru, and Achilonu 2024). This expansion likely conferred resilience against diverse stressors, including UV radiation, nutrient limitation, and fluctuations in salinity and temperature.

As a sulfur-containing compound, glutathione is closely linked to sulfur metabolism through DMSP and DHPS pathways (Noctor et al. 2012). Both metabolites play pivotal roles in diatom–bacteria interactions, serving as major sources of carbon and sulfur for marine bacteria. Our genomic analyses revealed that diatoms have increased copy numbers of genes for DHPS synthesis, but not of those involved in DMSP synthesis (**Supplementary Fig. S1, Fig. S2, Table S3**). This supports the emerging view that diatoms are key producers of DHPS in the ocean (Durham et al. 2015).

Our whole-genome analysis confirms that centric diatoms diversified prior to pennate diatoms, consistent with prior SSU rRNA studies (Medlin et al. 1996) (**Fig. 1**). Our study revealed expanded gene families related to microtubules and cellular protrusions in centric diatoms, likely enhancing their buoyancy (**Fig. 9**). In contrast, pennate diatoms exhibited bilateral symmetry and glide motility driven by actin-myosin-mediated raphe movement (Medlin et al. 1996; N. C. Poulsen et al. 1999). In pennate diatoms, genes related to actin, myosin, and cytoskeleton showed expansion, further underpinning their motility (**Fig. 9**) (Davutoglu et al. 2024). The expansion of distinct gene families in centric and pennate diatoms could reflect their adaptation to different ecological niches.

*N. inconspicua* showed significant gene expansion, without recent TE bursts or WGD. This expansion shows strong functional enrichment in core cellular processes (**Figs. S3, S4**), suggesting adaptive retention linked to its ecological niche. Future studies with chromosome-level assemblies and comparative analyses of closely related species would help clarify the mechanistic basis of this expansion.

### The impact of TEs on diatom evolution: evidence of TE diversity and insertion timing in diatom genomes

The TE content in the genomes of pennate diatoms has remained stable throughout evolution (below 30%), whereas centric diatoms exhibit greater variability, ranging from 5% to 50% (**Fig. 2**). The variation in TE proportions in centric diatoms partially reflects the variation in TE activity within their genomes, which may have different impacts on the stability of their genome structures. Our results demonstrate a relatively high proportion of LTR retrotransposons (LTR-RTs) in most diatom genomes, consistent with previous studies (Vitte and Panaud 2005). This conserved genomic feature mirrors the prevalence of LTR-RTs across diverse eukaryotic organisms, where these mobile elements are recognized as major drivers of structural and functional genome evolution (Tenaillon, Hollister, and Gaut 2010). Interestingly, the araphid pennate diatom *F. crotonensis* displayed a particularly remarkable LTR-RTs dominance, accounting for over 90% of its total TE content. This suggests that LTRs contribute significantly to genome architecture evolution in *F. crotonensis* (**Fig. 2**).

The raphe-lacking pennate araphid diatoms represent transitional forms in diatom evolution. Comparative genomic analysis showed that the pennate araphid diatoms *P. japonica* and *F. crotonensis*, diverged approximately 160.87 and 97 million years ago, respectively (**Fig. 1**). Our results suggest that TEs, particularly LTR-RTs, likely facilitated the evolutionary shift from centric to araphid pennate diatoms, as demonstrated by LTR diversity and unique LTR insertion timings. Firstly, the higher LTR-RTs diversity in *P. japonica* and *F. crotonensis* compared to other diatom species suggests increased potential for TE-mediated genomic plasticity (Casacuberta and González 2013) (**Fig. 3**). Secondly, LTR insertion time analysis revealed two distinct patterns of diatom genome reorganization: while most diatom genomes showed recent LTR insertion peaks, *P. japonica* and *F. crotonensis* displayed a unique pattern with a peak in a more ancient period (**Fig. 4**). This suggests that during early divergence, the genomes of these two araphid pennate diatoms underwent a major LTR transposon insertion event.

Our LTR/Copia and LTR/Gypsy insertion time analysis revealed that the vast majority of detectable LTR insertions across diatom genomes occurred within the last 2 million years, with only *P. japonica* retaining a substantial signature of ancient (3-5 Mya) insertions (**Fig. 5**). The predominance of recent insertions in most species indicates not a lack of ancient TE copies, but the progressive evolutionary elimination of old TE copies through illegitimate recombination, genome rearrangement, and purifying selection (Devos, Brown, and Bennetzen 2002). The retention of ancient LTRs in araphid pennates (*P. japonica* and *F. crotonensis*), potentially due to reduced deletion, selective pressure, or past massive proliferation, underscores the important role of historical TE activity in their early evolution (**Fig. 4**).

### The relationships of WGD, gene family dynamic, and TE insertion, and the hidden genomic mysteries

Parks et al. 2018 identified two deep WGD events occurring at ∼200 Mya and ∼170 Mya, which established an ancient polyploid ancestry for most extant diatoms. Our estimates of major lineage divergence times in diatoms (∼202 Mya and ∼173 Mya) are highly consistent with these timings (**Fig. 10**) and further reinforce the robustness of these events as major genomic landmarks. These two periods likely coincide with critical phases of early diatom diversification, when polyploidy may have facilitated genomic innovation, ecological expansion, and the emergence of lineage-specific traits.

**Fig. 10.**
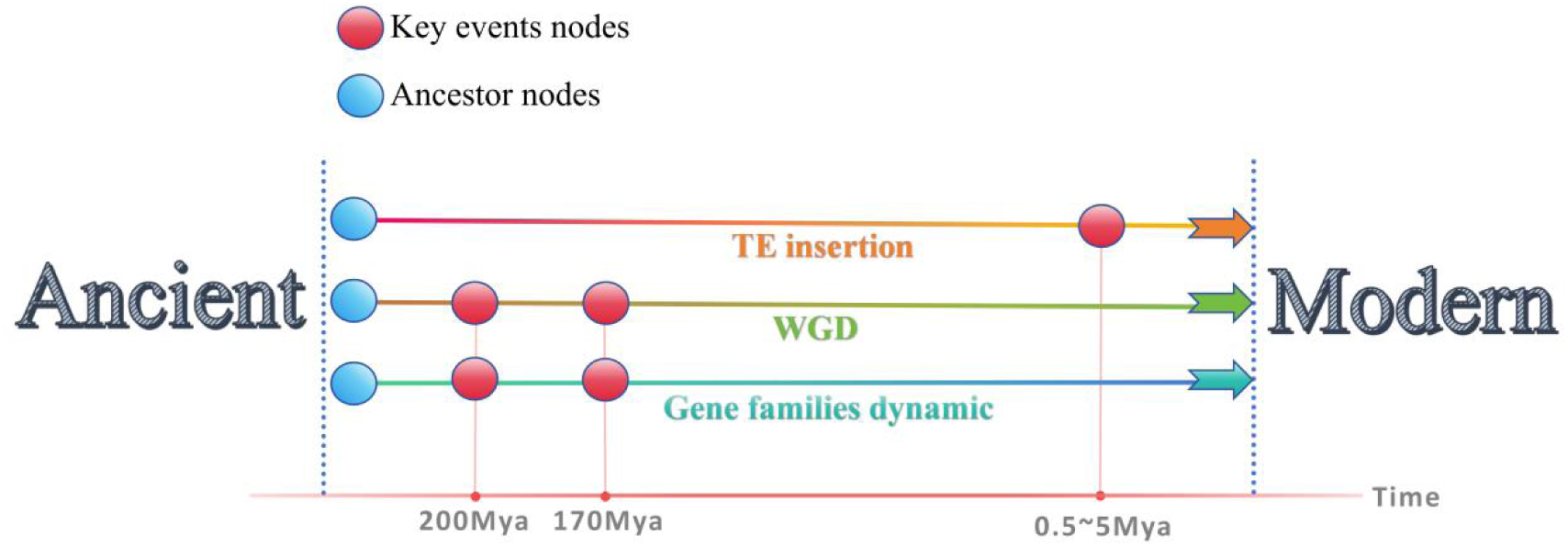
Temporal relationship between WGD, Gene family dynamics, and TE Insertion in diatoms. Two ancient WGD events (∼200 Ma and ∼170 Ma) coincide with divergence of major ancestral nodes and likely drove large-scale gene family expansions and contractions. TE insertions (0.5-5 Ma) occurred later than WGD and gene family dynamics, consistent with TE proliferation lagging behind WGD. The timeline is based on mutation rates (Krasovec et al. 2019) and reflects relative, not absolute, timing. Red nodes mark key genomic events; blue nodes indicate ancestral divergence points.

Building on previous findings that WGDs precede large-scale gene family reorganization (Gout et al. 2023; Zhang et al. 2025), we propose that large-scale gene family expansions in diatoms may reflect WGD-driven retention patterns, potentially linked to ecological adaptation and the diversification of diatoms. Our analysis added a functional dimension to this temporal framework by identifying the specific gene families that were significantly expanded or contracted, directly illustrating the lineage-specific genomic restructuring.

The temporal relationship between WGD and TE expansion has long been debated, but it is generally accepted that TE proliferation occurs after WGD. The “genome shock” hypothesis proposed that WGD triggers immediate bursts of TE activity (McClintock 1984). However, increasing evidence suggests that large-scale TE accumulation is typically not an abrupt burst but rather a gradual process. Empirical studies in *Atlantic salmon* further showed that some TE families became more active after WGD and contributed to cis-regulatory evolution (Monsen et al. 2025). TE proliferation has also been linked to rapid genome size increases, either independently or jointly with WGD, as reported in maize (Chandler and Brendel 2002) and *Corydoradinae catfishes* (Marburger et al. 2018).

Our LTR insertion time analysis revealed a striking temporal pattern: the vast majority of detectable LTR/Copia and LTR/Gypsy insertions across diatom genomes occurred within the last 0.5-5 million years (**Fig. 10**), substantially postdating both the ancient WGD events (∼200-170 Mya) and the major gene family turnover associated with these polyploidy events (**Fig. 10**). This apparent temporal gap between WGD (∼200-170 Mya) and observable TE proliferation (<5 Mya) is striking and requires careful interpretation. Critical methodological limitations compound these interpretive challenges. Our TE insertion time estimates assume a constant substitution rate from *P. tricornutum* (Krasovec et al. 2019), yet TE mutation rates vary substantially among taxa (Marchani et al. 2009). Moreover, our results readily detected recent TE insertions (<5 Mya) in modern diatom genomes, any hypothetical TE bursts coinciding with the ancient WGDs (100-200 Mya) likely have been largely erased by the pervasive loss of TE copies over time, thereby leaving minimal genomic signatures (Devos et al. 2002). The mutation rate uncertainty, combined with the progressive deletion of ancient TE copies, make it difficult to determine when the ancient TE insertions occurred.

Despite these limitations, our data provide the evidence in diatoms for distinct temporal scales of genome evolution following WGD: gene family reorganization likely occurred relatively rapidly, leaving stable phylogenetic signatures, whereas TE-mediated genome dynamics represent ongoing processes whose ancient history has likely been progressively obscured (**Fig. 10**). Future studies incorporating improved TE age dating methods and broader phylogenetic sampling will be essential to refine these temporal relationships and test the generality of these patterns across diatom lineages.

### Differences in functional genes and TEs among *P*. *tricornutum* strains

Gene families associated with DNA integration and histone modification have expanded in *P. tricornutum*, suggesting that DNA transposition and epigenetic regulation potentially play important roles in its environmental adaptability (**Fig. 9**). Our phylogenetic analysis identifies Pt4 and PtECS as the earliest-diverged and most recently derived strains of *P. tricornutum*, respectively (**Fig. 6**). PtECS has a unique cruciform structure that changes with temperature (He, Han, and Yu 2014). PtECS strain’s distinct cruciform shape appears associated with the expansion and contraction of specific gene families (**Fig. 6**). In PtECS, the expansion of gene families associated with protein tyrosine kinase and ankyrin repeats could aid cell signaling and differentiation, and maintain cellular integrity (Hunter 2000; Kumar and Balbach 2021) while the contraction of metallopeptidase-related gene families can influence the extracellular matrix and cell structure (Murphy and Nagase 2008). This association underscores the significance of gene family dynamics in shaping strain-specific morphologies.

The Pt4 strain, isolated from the low-salinity Baltic Sea, is adapted to low light and exhibits reduced non-photochemical quenching (Rastogi et al. 2020b). Consistent with its niche, we observed contraction in gene families involved in osmotic stress, light response, and organelle fission. Notably, while short-read sequencing revealed an increased copy number of the nitrate assimilation gene Phatr3_EG02286 in Pt4 (Rastogi et al. 2020b). Our long-read-based analysis further identified expansions in gene families related to amine metabolism (**Fig. 6**). These results indicate potential differences in nitrogen metabolism between Pt4 and other strains, warranting further investigation. Moreover, Pt4 exhibited the most significant reduction in dissolved inorganic carbon affinity under high CO₂ (1,000 ppm) among *P. tricorntum* strains, suggesting a unique carbon metabolism strategy (Huang et al. 2023). Pt4 exhibited an expansion of C4-related PPDK genes, whereas other strains had more C3-related genes (**Supplementary Table S4**), further supporting divergent carbon assimilation pathways across strains.

Our results demonstrate that TEs potentially drive divergence and adaptation in *P. tricornutum* strains. The widespread presence of LTR/Copia elements across all *P. tricornutum* strains suggests their key role in genome evolution (**Fig. 7A**). Notably, PtECS— identified as the most recently evolved strain (**Fig. 6**)—exhibits the highest LTR-RT content and the lowest TIR element abundance, consistent with the hypothesis that recent transposable element expansions drive genomic adaptation to environmental changes (Bourque et al. 2018). Strains from China (PtSCS and PtECS) share a higher number of common LTR/Copia elements and similar LTR insertion peaks during earlier periods, underscoring the role of LTR-RTs in *P. tricornutum* regional strain differentiation (**Fig. 7B**, **7C**).

Our study, utilizing long-read sequencing, significantly advances the characterization of TEs in *P. tricornutum*. While Maumus et al. 2009 first identified CoDi elements constituting ∼5.4% of the genome, subsequent work by Rastogi et al. 2018 revealed a higher proportion (∼7.6%). Our study extends these findings by discovering previously unclassified sequences that form a potentially novel CoDi branch (**Fig. 8**). Notably, the Chinese strains PtSCS and PtECS each contained nine identical novel sequences - a conserved pattern absents in other geographical variants. This distinct regional signature in transposon profiles suggests LTR/Copia retrotransposons contribute to geographical differentiation, potentially reflecting adaptive evolution to unique environmental pressures in Chinese coastal waters. The relatively higher retention of intact LTRs in PtECS may indicate either more recent transposition activity or reduced recombination rates in this lineage, preserving potentially autonomous elements that could retain transposition capacity (**Fig. 7D)**.

Our molecular dating suggests an unusually deep divergence (∼41 Mya) among *P. tricornutum* strains. While the absolute date requires caution, it is consistent with the known existence of four deeply diverged, globally distributed clades that exhibit significant genomic, morphological (oval, fusiform, and triradiate and cruciform), and physiological differentiation, indicating possible cryptic speciation (He et al. 2014; Rastogi et al. 2020a).

## CONCLUSION

Through long-read sequencing and comparative genomics, this study reveals the crucial roles of TEs and gene family dynamics in diatom evolutionary success. Comparative genomic analyses showed that diatom-specific expansions in polyamine metabolism and GST gene families underpin adaptive traits such as silica wall formation and oxidative stress resistance in diatoms. Divergent gene family profiles between centric and pennate diatoms correlate with their distinct morphologies and motility strategies, while TE insertion timing suggests TEs played important roles in the evolutionary transition to raphid pennate diatoms.

Strain-level analyses of *P. tricornutum* reveal geographic adaptation signatures, with morphology-related gene expansions in East China Sea isolates and metabolic gene contractions in Baltic Sea isolates. TE diversity correlates with geographic distribution, demonstrating how TE-mediated genome plasticity drives local adaptation.

Our phylogenomic analyses reveal temporal consistency between major lineage divergences (∼202 and ∼173 Mya) and ancient whole-genome duplications (∼200 and ∼170 Mya). Gene family analyses reveal lineage-specific expansions and contractions at these divergence nodes. TE expansions show more recent timescales, reflecting progressive deletion of ancient copies while ongoing accumulation generates the recent insertion peaks observed across diatom lineages.

## DATA AVAILABILITY

The sequencing data have been uploaded to NCBI (project ID: PRJNA1112327).

## ACKNOWLEDGEMENTS

This study was supported by the National Natural Science Foundation of China (42076128) and

Natural Science Foundation of Fujian Province of China (2024J01020)

## AUTHOR CONTRIBUTIONS

Xin Lin conceived and designed the experiments. Xu Zhang performed the experiments. Tianze Zheng and Xu Zhang analyzed data. Tianze Zheng, Tianren Liu and Xinzhu Liu wrote the manuscript. Chris Bowler reviewed the manuscript and provided revision suggestions. Xin Lin revised the manuscript. All authors reviewed the manuscript.

## CONFLICT OF INTEREST STATEMENT

The authors declare that they have no conflicts of interest related to this work.

## ANIMAL AND HUMAN RIGHTS STATEMENT

This study did not involve any human participants or animals. Therefore, no ethical approval was required.

## Notes

### Competing Interest Statement

The authors have declared no competing interest.

