## Supplementary material for "The roles of Transposable Elements and Gene Family Dynamics in Shaping Diversity and Evolution in Diatoms": Supplymentary File

**Supplementary materials**

As detailed by Durham et al.(Durham et al. 2019), diatoms are among the most diversified producers of sulfur metabolites, capable of producing compounds like DMSP (dimethylsulfoniopropionate), DHPS (dehydropantetheine phosphate), and taurine. DMSP and DHPS are significant chemicals involved in the marine sulfur cycle. They are primarily produced by phytoplankton and utilized by marine bacteria as a carbon source, serving as key materials exchanged between bacteria and algae. Phytoplankton mainly synthesize DMSP through the transamination pathway of methionine. The critical step in this pathway has been identified as the third step, the methylation process. The key gene involved in DMSP synthesis in *T. pseudonana*, TpMMT, has been discovered. Comparative analysis based on this sequence has revealed the expansion and contraction of the methyltransferase family in diatoms. Research on DHPS is not as extensive as DMSP. However, the synthesis pathway for DHPS has been relatively well elucidated.

Therefore, we focused on the copy number of the key enzyme genes involved in the synthesis of DMSP (Tpmmt, Dsyb), DHPS synthase, and glutathione S-transferases in the diatom genome (**Table S3**). Comparisons showed that Tpmmt (a key gene for DMSP synthesis identified in diatoms) has a certain number of copies in most diatoms, while Dsyb (a key gene for DMSP synthesis found in protozoa) was found to have one copy each in *F. cylindrus*, *S. robusta*, and *S. marinoi*. The enzymes for DHPS synthesis, SDH, and CoA, have a certain number of copies in all diatoms. For comC, except in species like *C. tenuissimus*, *F. cylindrus*, *S. robusta*, *T.* *pseudonana*, *S.* *marinoi*, and *C.* *cryptica*, the rest of the species have a certain number of copies, with *N.* *inconspicua* having the highest number at two copies, while the rest have one.

An increase in the number of copies of PYC1 (pyruvate carboxylase) was observed in Pt1. In addition, PtSLC4 and TpSLC4 genes in PT1 and PTSCS displayed variations and potential enhancements in the C3 pathway. Notably, the significant expansion of the PPDK gene family and its high copy number in the Pt4 genome may reflect a unique response to high carbon environments, consistent with earlier studies(Huang et al. 2020, 2023) (**Table S4**) .

**Research pipeline of genome comparison of different *P. tricornutum* strains**

To investigate genomic diversity among *P. tricornutum* strains, we performed multi-scale comparative analyses encompassing structural variation, gene family dynamics, and transposable element characterization. Assembly quality metrics including contig N50, total length, GC content were calculated using Assembly-stats. Structural variations (SVs) including insertions, deletions, duplications, and inversions were detected from long-read sequencing data using Sniffles (Sedlazeck et al. 2018). Comparative genomic analyses, including divergence time estimation and gene family expansion/contraction analysis, were conducted following the pipeline described above (see “Comparative genomic analysis of diatoms and selected coccolithophore, red and green algal species”). To examine variation in CCM pathway components, we identified genes associated with C_3_ and C_4_ carbon fixation pathways across P. tricornutum strains using BLASTP (E-value < 1e-5, identity > 80%) with known CCM genes as queries. TE annotation and dating analyses were conducted following the pipeline described above (see “TE annotation and divergence analysis of diatoms and representative red algal and green algal species”).

Given the demonstrated importance of LTR/Copia elements, particularly CoDi elements, in P. tricornutum genome evolution (Maumus et al. 2009), we performed detailed characterization of this superfamily. Reverse transcriptase (RT) domains were extracted from full-length LTR elements and clustered to identify shared and strain-specific LTR/Copia families across the four strains. Phylogenetic relationships among RT domains were reconstructed using RAxML.

Regarding the analysis of Intact/Solo LTRs in different strains of *P. tricornutum*, we primarily organized and summarized the annotation results using a custom R script.

**Table S1.** transposon contents of different algae

| Species | Class Ⅰ | | | Class Ⅱ | | | Unkown | Total |
| --- | --- | --- | --- | --- | --- | --- | --- | --- |
|  | LTR | LINE | Others | TIR | Helitron | Others |  |  |
| *Phaeodactylum tricornutum 1*(Filloramo et al. 2021) | 12% | 0.04% | 0.00% | 3.21% | 0.31% | 0.00% | 1.64% | 17.20% |
| *Phaeodactylum tricornutum (Pt4)* | 12.83% | 0.00% | 0.04% | 3.16% | 0.42% | 0.00% | 1.14% | 17.72% |
| *Phaeodactylum tricornutum (PtSCS)* | 12.11% | 0.04% | 0.00% | 2.11% | 0.42% | 0.00% | 0.89% | 15.60% |
| *Phaeodactylum tricornutum (PtECS)* | 12.9% | 0.00% | 0.06% | 1.97% | 0.4% | 0.03% | 0.76% | 16.11% |
| *Cyclotella cryptica*(Nelson et al. 2021) | 34.98% | 0.64% | 0.29% | 6.8% | 1.76% | 0.13% | 8.75% | 53.37% |
| *Fragilariopsis cylindrus*(Paajanen et al. 2017) | 5.35% | 0.49% | 0.35% | 3.47% | 0.51% | 0.00% | 2.25% | 12.43% |
| *Fistulifera pelliculosa* | 4.46% | 0.23% | 0.04% | 2.58% | 0.33% | 0.00% | 1.36% | 8.99% |
| *Minidiscus variabilis* | 4.46% | 0.20% | 0.00% | 3.79% | 1.19% | 0.00% | 2.12% | 11.78% |
| *Nitzschia inconspicua*(Oliver et al. 2021) | 8.09% | 0.62% | 0.42% | 5.09% | 1.61% | 0.02% | 5.96% | 21.82% |
| *Nitzschia putrida*(Nelson et al. 2021) | 5.03% | 0.06% | 0.00% | 3.31% | 0.14% | 0.00% | 2.48% | 12.67% |
| *Psammoneis japonica* | 12.02% | 0.24% | 0.6% | 8.36% | 1.09% | 0.00% | 4.69% | 26.98% |
| *Skeletonema costatum*(Sorokina et al. 2022) | 8.42% | 0.21% | 0.04% | 2.45% | 1.01% | 0.00% | 5.87% | 18.00% |
| *Seminavis robusta*(Osuna-Cruz et al. 2020) | 15.74% | 0.89% | 0.35% | 1.87% | 0.95% | 0.00% | 3.44% | 23.24% |
| *Thalassiosira oceanica*(Nelson et al. 2021) | 8.19% | 0.14% | 0.24% | 6.98% | 0.08% | 0.00% | 10.76% | 26.40% |
| *Chaetoceros muellerii*(Nelson et al. 2021) | 12.37% | 0.36% | 0.94% | 2.94% | 0.48% | 0.07% | 4.35% | 21.51% |
| *Chaetoceros calcitrans*(Nelson et al. 2021) | 2.08% | 0.02% | 0.04% | 1.92% | 0.13% | 0.00% | 1.23% | 5.43% |
| *Chaetoceros neogracile*(Nelson et al. 2021) | 1.92% | 0.00% | 0.06% | 2.77% | 0.6% | 0.00% | 2.02% | 7.37% |
| *Amphora coffeaeformis*(Nelson et al. 2021) | 1.28% | 0.05% | 0.02% | 1.32% | 0.18% | 0.00% | 0.36% | 3.22% |
| *Skeletonema marinoi*(Nelson et al. 2021) | 2.41% | 0.08% | 0.02% | 5.41% | 0.22% | 0.00% | 3.67% | 11.81% |
| *Cylindrotheca fusiformis*(Nelson et al. 2021) | 5.48% | 0.37% | 0.59% | 5.68% | 1.51% | 0.00% | 4.30% | 17.94% |
| *Skeletonema dohrnii*(Nelson et al. 2021) | 2.9% | 0.01% | 0.05% | 6.92% | 1.05% | 0.00% | 4.09% | 15.04% |
| *Skeletonema menzelii*(Nelson et al. 2021) | 1.99% | 0.00% | 0.00% | 2.79% | 0.05% | 0.00% | 2.75% | 7.59% |
| *Attheya sp.*(Nelson et al. 2021) | 11.23% | 0.26% | 0.3% | 15.84% | 0.07% | 0.01% | 20.13% | 47.84% |
| *Thalassiosira pseudonana*(Filloramo et al. 2021) | 3.53% | 0.30% | 0.01% | 2.46% | 0.56% | 0.00% | 1.38% | 8.24% |
| *Navicula incerta*(Nelson et al. 2021) | 11.95% | 0.41% | 0.46% | 4.36% | 0.07% | 0.00% | 3.31% | 20.72% |
| *Navicula pelliculosa*(Nelson et al. 2021) | 0.91% | 0.27% | 0.28% | 5.67% | 0.23% | 0.00% | 1.75% | 9.12% |
| *Cyclotella meneghiniana*(Nelson et al. 2021) | 2.35% | 0.25% | 0.27% | 9.48% | 0.06% | 0.05% | 28.27% | 40.74% |
| *Mayamaea pseudoterrestris*(Suzuki et al. 2022) | 14.17% | 0.00% | 0.00% | 2.13% | 0.16% | 0.00% | 0.00% | 16.46% |
| *Epithemia pelagica* | 13.11% | 0.00% | 0.00% | 0.00% | 0.52% | 0.00% | 0.00% | 13.64% |
| *Fragilaria crotonensis*(Zepernick et al. 2022) | 13.66% | 0.00% | 0.00% | 0.01% | 0.30% | 0.00% | 0.00% | 13.98% |
| *Craspedostauros australis* | 17.95% | 0.00% | 0.00% | 0.00% | 1.18% | 0.00% | 0.00% | 19.13% |
| *Chaetoceros tenuissimus*(Hongo et al. 2021) | 5.27% | 0.15% | 0.13% | 3.09% | 0.37% | 0.00% | 2.09% | 11.10% |
| *Chlorella vulgaris*(Cecchin et al. 2019) | 2.33% | 0% | 0.00% | 2.12% | 0.29% | 0.00% | 0.00% | 4.75% |
| *Scenedesmus sp.*(Calixto Mancipe, McLaughlin, and Barney 2022) | 1.35% | 0% | 0.00% | 0% | 0.05% | 0% | 0% | 1.40% |
| *Volvox africanus*(Yamamoto et al. 2021) | 10.09% | 0% | 0% | 0% | 1.24% | 0% | 0% | 13.44% |
| *Galdieria partita*(Hirooka et al. 2022) | 0% | 0% | 0% | 2.09% | 0.15% | 0% | 0% | 2.24% |
| *Porphyridium purpureum*(Lee et al. 2019) | 7.59% | 0% | 0% | 0.72% | 0.20% | 0% | 0% | 8.52% |
| *Gracilariopsis chorda*(Lee et al. 2018) | 35.37% | 0% | 0% | 4.99% | 2.33% | 0% | 0% | 42.69% |
| *Porphyra umbilicalis*(Brawley et al. 2017) | 14.15% | 0% | 0% | 8.88% | 1.31% | 0% | 0% | 24.33% |
| *Pyropia yezoensis*(Zhang et al. 2023) | 29.92% | 0% | 0% | 10.13% | 0% | 0% | 5.87% | 47.07% |

**Table S2.** Gene Enrichment Analysis for Contraction and Expansion

| Gene family dynamics | Eenrichment analysis forms | Enrichment result | Illustrate | Classification |
| --- | --- | --- | --- | --- |
| Expansion | GO-BP | Carbohydrate metabolic process | - | Diatoms |
|  |  | Nitrogen cycle metabolic process |  |  |
|  |  | Microtubule-based movement |  |  |
|  |  | Tetraterpenoid metabolic process | Metabolic process combining pigment and antioxidant functions |  |
|  |  | Carotenoid metabolic process |  |  |
|  |  | Regulation of phosphorus metabolic process | - |  |
|  |  | Protein peptidyl-prolyl isomerization | Processes related to protein structure |  |
|  |  | Regulation of protein modification process |  |  |
|  |  | Protein ubiquitination |  |  |
|  |  | Response to oxidative stress | - |  |
|  | KEGG | Alcoholism | Related to histone modification |  |
|  |  | Carbon fixation in photosynthetic organisms | Related to carbon metabolism |  |
|  |  | Carbon metabolism |  |  |
|  |  | Methane metabolism | Related to formaldehyde metabolism |  |
|  |  | Glutathione metabolism | Discovery of genes related to Glutathione S-transferase and Glutathione peroxidase |  |
| Contraction | GO-BP | Phosphorylation | Phosphorylation-related processes |  |
|  |  | Protein phosphorylation |  |  |
|  | KEGG | Oxytocin signaling pathway | Multiple signaling pathways |  |
|  |  | Phospholipase D signaling pathway |  |  |
|  |  | Adipocytokine signaling pathway |  |  |
|  |  | Rap1 signaling pathway |  |  |
|  |  | AMPK signaling pathway |  |  |
|  |  | Insulin signaling pathway |  |  |
|  |  | Apelin signaling pathway |  |  |
|  |  | Focal adhesion | Pathways related to cytoskeleton and cell junctions |  |
|  |  | Regulation of actin cytoskeleton |  |  |
|  |  | Tight junction |  |  |
| Expansion | GO-BP | DNA integration | Gene recombination or transposition related processes | Pennates |
|  |  | DNA metabolic process |  |  |
|  |  | Nitrogen cycle metabolic process | Genes found to be related to nitrite transport and nitrate transmembrane transport |  |
|  | GO-CC | myosin complex | Actin, myosin, cytoskeleton related |  |
|  |  | actin cytoskeleton |  |  |
|  |  | cytoskeleton |  |  |
|  | KEGG | Phenylalanine metabolism | - |  |
|  |  | Phenylalanine, tyrosine and tryptophan biosynthesis |  |  |
|  |  | Glutathione metabolism |  |  |
|  |  | Nitrogen metabolism |  |  |
| Contraction | GO-BP | DNA conformation change |  |  |
|  |  | Microtubule-based process |  |  |
|  |  | Phosphorylation |  |  |
|  | KEGG | Alcoholism |  |  |
|  |  | Systemic lupus erythematosus |  |  |
|  |  | Viral carcinogenesis |  |  |
|  |  | Cellular senescence |  |  |
|  |  | Gap junction |  |  |
| Expansion | GO-BP | microtubule-based movement | movement and transport of the cytoskeleton (microtubules) |  |
|  |  | microtubule-based process | Processes and structures related to microtubules and the cytoskeleton | Centrales |
|  |  | transport along microtubule |  |  |
|  |  | microtubule-based transport |  |  |
|  | GO-CC | Dynein complex |  |  |
|  |  | Microtubule associated complex |  |  |
|  |  | Microtubule |  |  |
|  |  | Polymeric cytoskeletal fiber |  |  |
|  |  | Supramolecular polymer |  |  |
|  |  | Microtubule cytoskeleton |  |  |
|  |  | Axoneme |  |  |
|  |  | Cytoskeleton |  |  |
|  |  | Cilium | Related to the movement characteristics of centric diatoms |  |
|  |  | Cell projection |  |  |
|  | KEGG | Glutathione metabolism | - |  |
|  |  | Alcoholism |  |  |
| Contraction | GO-BP | glutamine family amino acid biosynthetic process |  |  |
|  |  | carboxylic acid biosynthetic process |  |  |
|  |  | organic acid biosynthetic process |  |  |
|  |  | glutamine family amino acid metabolic process |  |  |
|  |  | arginine metabolic process |  |  |
|  |  | vesicle budding from membrane |  |  |
| Expansion | GO-BP | DNA Integration | DNA metabolism and integration | *P. tricornutum* |
|  |  | DNA Metabolic Process |  |  |
|  |  | Histone Lysine Methylation | Histone methylation modification related |  |
|  |  | Histone Methylation |  |  |
|  |  | Peptidyl-Lysine Methylation |  |  |
|  |  | Protein Methylation |  |  |
|  |  | Macromolecule Methylation |  |  |
|  |  | Histone modification |  |  |
|  |  | Peptidyl-lysine modification |  |  |
|  |  | Protein Alkylation | Chemical modification and degradation processes targeting proteins |  |
|  |  | Proteolysis |  |  |
|  |  | Cellular Response to Oxidative Stress | Cellular Response to Stress |  |
|  |  | Cellular Response to Chemical Stress |  |  |
|  |  | Regulation of hydrolase activity | Regulation of various enzymatic activities |  |
|  |  | positive regulation of catalytic activity |  |  |
|  |  | positive regulation of molecular function |  |  |
|  | GO-CC | nucleoplasm | - |  |
|  |  | Histone Methyltransferase Activity | Histone methylation modification related |  |
|  |  | Protein-Lysine N-Methyltransferase Activity |  |  |
|  |  | Lysine N-Methyltransferase Activity |  |  |
|  |  | Protein Methyltransferase Activity |  |  |
|  |  | Histone Lysine N-Methyltransferase Activity |  |  |
|  |  | Serine-Type Endopeptidase Activity | Serine protease and hydrolase |  |
|  |  | Serine-Type Peptidase Activity |  |  |
|  |  | Serine Hydrolase Activity |  |  |
|  | KEGG | Lysine Degradation | - |  |
|  |  | Arginine and Proline Metabolism |  |  |
|  |  | Ferroptosis |  |  |
|  |  | Pentose Phosphate Pathway |  |  |
|  |  | Carbon Fixation in Photosynthetic Organisms |  |  |
| Contraction | GO-BP | Microtubule-based Movement | Microtubule-associated |  |
|  |  | Microtubule-based Process |  |  |
|  |  | Response to Abiotic Stimulus |  |  |
|  |  | Proteolysis | - |  |
|  | KEGG | Protein processing in endoplasmic reticulum |  |  |
|  |  | Systemic lupus erythematosus |  |  |
|  |  | MicroRNAs in cancer |  |  |
|  |  | Gastric cancer |  |  |
|  |  | Legionellosis |  |  |

**Table S3.** Statistics on the copy number of genes in the sulfur metabolism and other pathway

|  | **DMSP** | | **DHPS** | | | | **Glutathione S-transferase** | **ChlC** | **CRTISO5** |
| --- | --- | --- | --- | --- | --- | --- | --- | --- | --- |
|  | **TpMMT** | **Dsyb** | **SDH** | **CoA23871** | **CoA42458** | **comC** |  |  |  |
| ***Chaetoceros tenuissimus*** | **0** | **0** | **2** | **2** | **1** | **0** | **3** | **2** | **5** |
| ***Fragilariopsis cylindrus*** | **1** | **1** | **3** | **2** | **1** | **0** | **2** | **1** | **7** |
| ***Fragilaria crotonensis*** | **0** | **0** | **5** | **3** | **3** | **1** | **2** | **1** | **5** |
| ***Nitzschia inconspicua*** | **3** | **0** | **7** | **6** | **2** | **2** | **4** | **3** | **9** |
| ***PT1*** | **2** | **0** | **1** | **2** | **1** | **1** | **2** | **1** | **7** |
| ***PT4*** | **2** | **0** | **2** | **2** | **1** | **1** | **2** | **1** | **5** |
| ***PT11*** | **2** | **0** | **1** | **2** | **1** | **1** | **2** | **1** | **4** |
| ***PT12*** | **2** | **0** | **1** | **2** | **1** | **1** | **2** | **1** | **5** |
| ***Seminavis robusta*** | **2** | **1** | **3** | **3** | **1** | **0** | **4** | **1** | **8** |
| ***Thalassiosira pseudonana*** | **2** | **0** | **3** | **2** | **1** | **0** | **2** | **1** | **5** |
| ***Mayamaea pseudoterrestris*** | **0** | **0** | **3** | **2** | **1** | **1** | **2** | **1** | **4** |
| ***Skeletonema marinoi*** | **4** | **1** | **5** | **3** | **1** | **0** | **2** | **1** | **6** |
| ***Cyclotella cryptica*** | **2** | **0** | **5** | **5** | **1** | **0** | **3** | **1** | **4** |
| ***Psammoneis japonica*** | **0** | **0** | **2** | **2** | **2** | **1** | **4** | **1** | **4** |

**Table S4.** Copy numbers of CCM genes related to C4 and C3 pathways in four strains of *P. tricornutum*

|  | **C4** | | | | **C3** | | | | | | |
| --- | --- | --- | --- | --- | --- | --- | --- | --- | --- | --- | --- |
|  | **PEPCK** | **PYC1** | **PYC2** | **PPDK** | **PtCA2** | **PtSLC4-2** | **PtSLC4-3** | **PtSLC4-5** | **PtSLC4-6** | **PtSLC26** | **TpSLC4-3** |
| **Pt1** | **1** | **6** | **5** | **1** | **2** | **6** | **7** | **7** | **4** | **1** | **5** |
| **Pt4** | **1** | **6** | **5** | **2** | **2** | **4** | **6** | **6** | **4** | **1** | **4** |
| **PtECS** | **1** | **5** | **5** | **1** | **1** | **5** | **7** | **7** | **4** | **1** | **4** |
| **PtSCS** | **1** | **5** | **5** | **1** | **2** | **4** | **6** | **6** | **4** | **1** | **4** |

**Table S5.** Gene function table of phytoplankton

| **Gene Name** | **Functional Category** | **Function Description** |
| --- | --- | --- |
| Fucoxanthin Chlorophyll a/c Binding Protein | Photosynthesis | Involved in the binding of pigments related to chlorophyll and algal photosynthesis. |
| Heat Shock Protein 70 | Stress Response | Helps cells cope with environmental stress like temperature and salinity by protecting protein folding. |
| Carbonic Anhydrase | Photosynthesis | Involved in the conversion of CO2, supporting carbon fixation during photosynthesis. |
| Ribulose-bisphosphate Carboxylase | Photosynthesis | A key enzyme in carbon fixation during photosynthesis. |
| β-1,3-glucanase | Cell Structure | Breaks down β-1,3-glucans, involved in cell wall synthesis and repair. |
| Procollagen-Proline 4-Dioxygenase Activity | Stress Response | Eliminates peroxides in cells, helping with oxidative stress. |

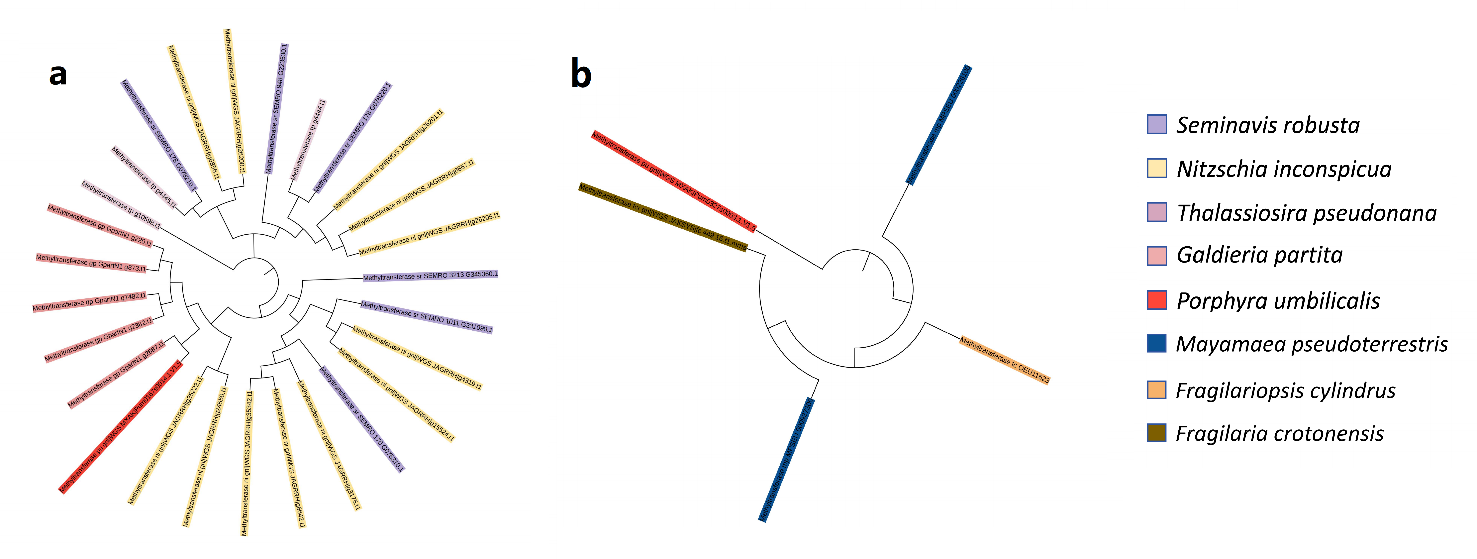

**Figure S1.** Phylogenetic Tree of Expanded and Contracted DMSP Key Enzyme Gene Families in Different Species. a.Expansion b.constraction

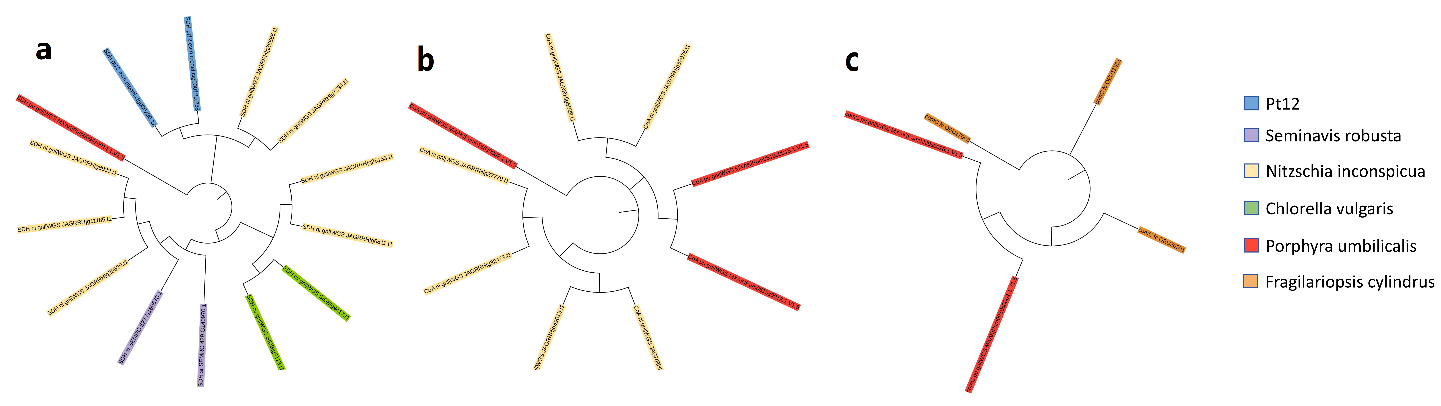

**Figure S2.** Phylogenetic Tree of Expanded DHPS Synthase Gene Families in Different SpeciesFigure.a.SDH b.CoA c.ComC

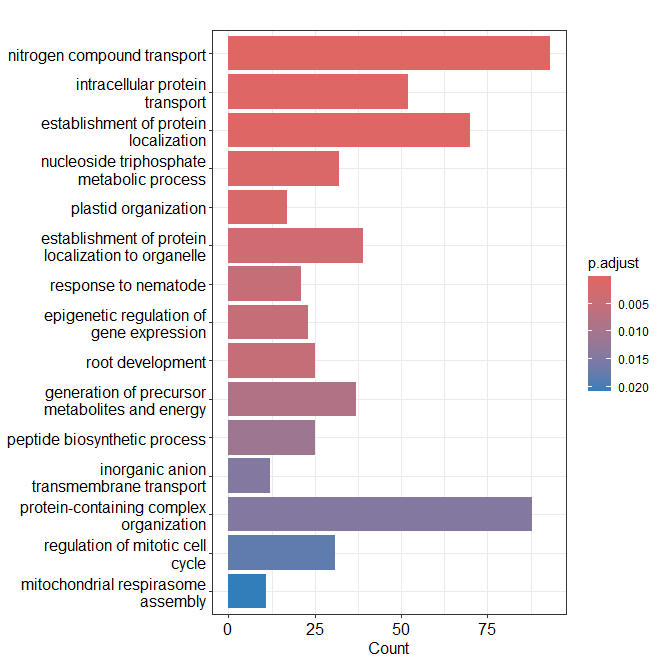

**Figure S3.** GO enrichment analysis of expanded gene families in N. inconspicua

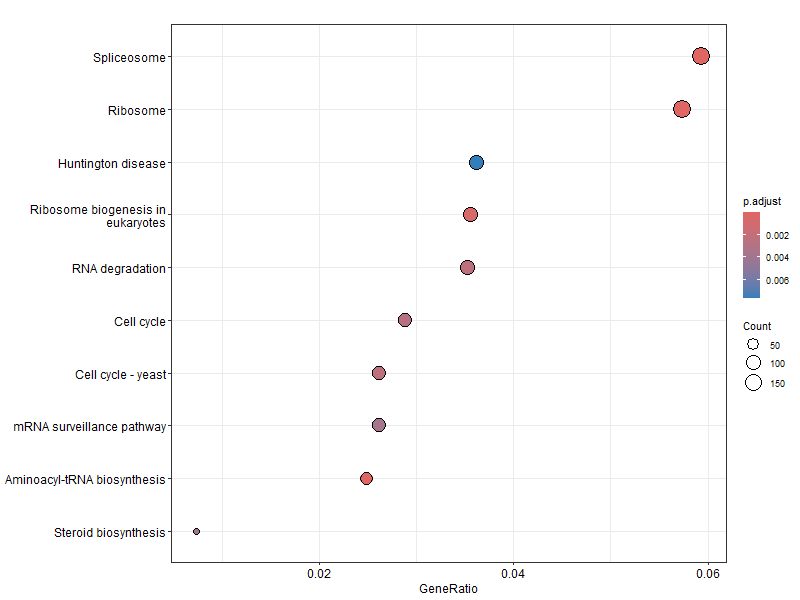

**Figure S4.** KEGG enrichment analysis of expanded gene families in N. inconspicua

**Whole-genome duplication analysis of araphid pennate diatoms**

To assess the potential contribution of whole-genome duplication (WGD) to diatom evolution, we conducted an exploratory analysis focusing on non-raphe pennate lineages, which occupy a transitional evolutionary position. Ks (synonymous substitution rate) distributions were calculated for paralogous gene pairs to identify potential duplication peaks (Fig. S5).

Ks analysis revealed prominent peaks at high Ks values (3.5–4.5) across all examined diatom species combinations, which could potentially reflect ancient duplication events in early diatom evolution. The *F. crotonensis*–*T. pseudonana* comparison showed a secondary peak at Ks=2.0–2.5, while other comparisons (*F. cylindrus*–*F. crotonensis*, *F. cylindrus*–*T. pseudonana*) showed primary peaks around Ks=4.0. Comparisons between *F. crotonensis* (araphid pennate) and *F. cylindrus* (raphid pennate) showed peaks at Ks=4.0–4.5, while the *F. cylindrus*–*T. pseudonana* (centric diatom) comparison exhibited even higher Ks values, suggesting progressive accumulation of genomic divergence across these lineages.

We acknowledge that the shapes of these Ks curves are complex and difficult to interpret definitively. It remains unclear whether genuine WGD signals can be reliably distinguished from other duplication mechanisms—such as large-scale segmental duplications, tandem gene duplications, or lineage-specific gene family expansions—based on Ks distributions alone. Ancient WGD events may also be obscured by saturation effects at high Ks values and by the complex evolutionary history of diatoms, including secondary endosymbiosis and substantial gene loss. Additional lines of evidence, including synteny analysis, chromosomal collinearity assessment, and broader taxonomic sampling, would be needed for robust conclusions. These results are therefore presented as preliminary patterns that warrant further investigation rather than definitive evidence for specific WGD events.

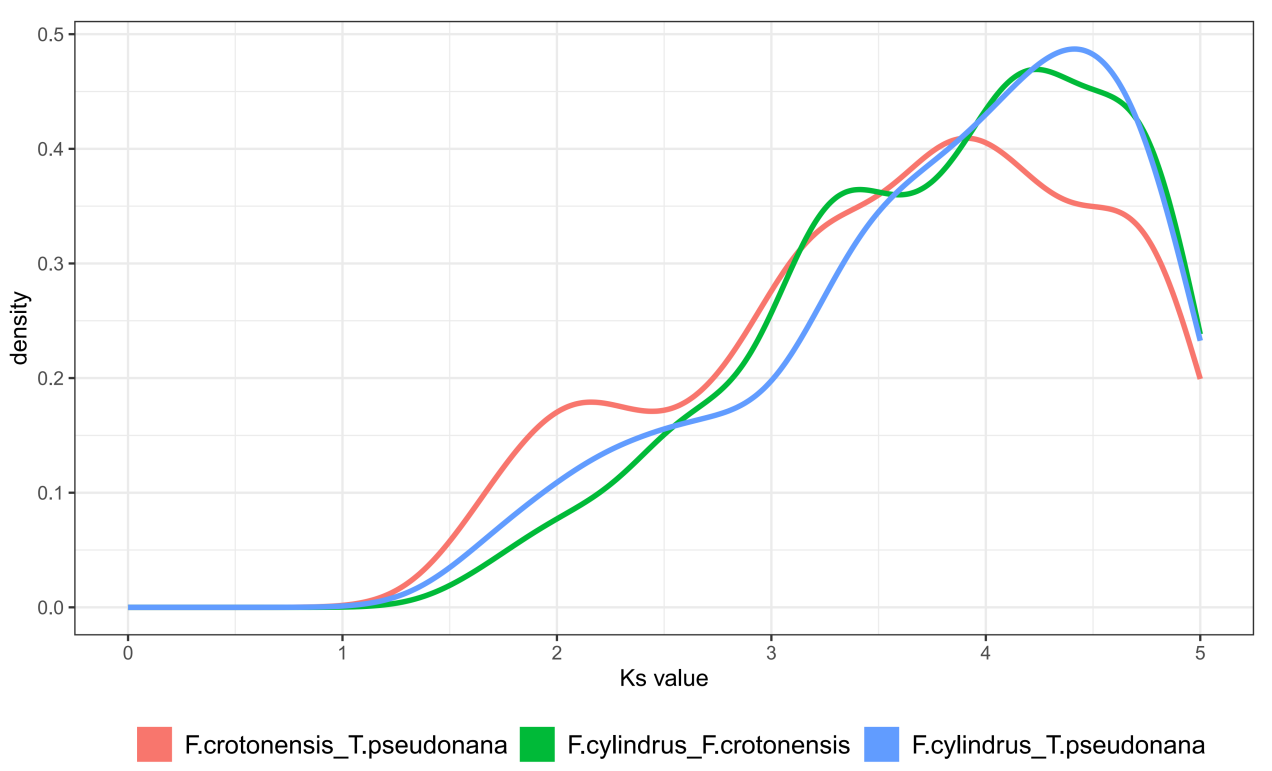

**Fig. S5 Whole-genome duplication events in *F. crotonensis*, *F. cylindrus*, and *T. pseudonana*. The x-axis (Ks value) represents the synonymous substitution rate, used to estimate the divergence time of genes, with higher values indicating earlier divergence; the y-axis (density) shows the density distribution of gene pairs. The peaks in the curve represent potential WGD events, with the position of each peak reflecting the relative timing of the duplication event. Different colored curves represent comparisons between different species combinations.**

**References：**

Brawley, Susan H., Nicolas A. Blouin, Elizabeth Ficko-Blean, Glen L. Wheeler, Martin Lohr, Holly V. Goodson, Jerry W. Jenkins, Crysten E. Blaby-Haas, Katherine E. Helliwell, Cheong Xin Chan, Tara N. Marriage, Debashish Bhattacharya, Anita S. Klein, Yacine Badis, Juliet Brodie, Yuanyu Cao, Jonas Collén, Simon M. Dittami, Claire M. M. Gachon, Beverley R. Green, Steven J. Karpowicz, Jay W. Kim, Ulrich Johan Kudahl, Senjie Lin, Gurvan Michel, Maria Mittag, Bradley J. S. C. Olson, Jasmyn L. Pangilinan, Yi Peng, Huan Qiu, Shengqiang Shu, John T. Singer, Alison G. Smith, Brittany N. Sprecher, Volker Wagner, Wenfei Wang, Zhi-Yong Wang, Juying Yan, Charles Yarish, Simone Zäuner-Riek, Yunyun Zhuang, Yong Zou, Erika A. Lindquist, Jane Grimwood, Kerrie W. Barry, Daniel S. Rokhsar, Jeremy Schmutz, John W. Stiller, Arthur R. Grossman, and Simon E. Prochnik. 2017. ‘Insights into the Red Algae and Eukaryotic Evolution from the Genome of Porphyra Umbilicalis (Bangiophyceae, Rhodophyta)’. *Proceedings of the National Academy of Sciences of the United States of America* 114(31):E6361–70. doi: 10.1073/pnas.1703088114.

Calixto Mancipe, Natalia, Evelyn M. McLaughlin, and Brett M. Barney. 2022. ‘Genomic Analysis and Characterization of Scenedesmus Glucoliberatum PABB004: An Unconventional Sugar-Secreting Green Alga’. *Journal of Applied Microbiology* 132(3):2004–19. doi: 10.1111/jam.15311.

Cecchin, Michela, Luca Marcolungo, Marzia Rossato, Laura Girolomoni, Emanuela Cosentino, Stephan Cuine, Yonghua Li-Beisson, Massimo Delledonne, and Matteo Ballottari. 2019. ‘Chlorella Vulgaris Genome Assembly and Annotation Reveals the Molecular Basis for Metabolic Acclimation to High Light Conditions’. *The Plant Journal: For Cell and Molecular Biology* 100(6):1289–1305. doi: 10.1111/tpj.14508.

Durham, Bryndan P., Angela K. Boysen, Laura T. Carlson, Ryan D. Groussman, Katherine R. Heal, Kelsy R. Cain, Rhonda L. Morales, Sacha N. Coesel, Robert M. Morris, Anitra E. Ingalls, and E. Virginia Armbrust. 2019. ‘Sulfonate-Based Networks between Eukaryotic Phytoplankton and Heterotrophic Bacteria in the Surface Ocean’. *Nature Microbiology* 4(10):1706–15. doi: 10.1038/s41564-019-0507-5.

Filloramo, Gina V., Bruce A. Curtis, Emma Blanche, and John M. Archibald. 2021. ‘Re-Examination of Two Diatom Reference Genomes Using Long-Read Sequencing’. *BMC Genomics* 22(1):379. doi: 10.1186/s12864-021-07666-3.

Hirooka, Shunsuke, Takeshi Itabashi, Takako M. Ichinose, Ryo Onuma, Takayuki Fujiwara, Shota Yamashita, Lin Wei Jong, Reiko Tomita, Atsuko H. Iwane, and Shin-Ya Miyagishima. 2022. ‘Life Cycle and Functional Genomics of the Unicellular Red Alga Galdieria for Elucidating Algal and Plant Evolution and Industrial Use’. *Proceedings of the National Academy of Sciences of the United States of America* 119(41):e2210665119. doi: 10.1073/pnas.2210665119.

Hongo, Yuki, Kei Kimura, Yoshihiro Takaki, Yukari Yoshida, Shuichiro Baba, Genta Kobayashi, Keizo Nagasaki, Takeshi Hano, and Yuji Tomaru. 2021. ‘The Genome of the Diatom Chaetoceros Tenuissimus Carries an Ancient Integrated Fragment of an Extant Virus’. *Scientific Reports* 11:22877. doi: 10.1038/s41598-021-00565-3.

Huang, Ruiping, Jiancheng Ding, Jiazhen Sun, Yang Tian, Chris Bowler, Xin Lin, and Kunshan Gao. 2020. ‘Physiological and Molecular Responses to Ocean Acidification among Strains of a Model Diatom’. *Limnology and Oceanography* 65(12):2926–36. doi: 10.1002/lno.11565.

Huang, Teng, Fan Hu, Yufang Pan, Chenjie Li, and Hanhua Hu. 2023. ‘Pyruvate Orthophosphate Dikinase Is Required for the Acclimation to High Bicarbonate Concentrations in Phaeodactylum Tricornutum’. *Algal Research* 72:103131. doi: 10.1016/j.algal.2023.103131.

Lee, JunMo, Dongseok Kim, Debashish Bhattacharya, and Hwan Su Yoon. 2019. ‘Expansion of Phycobilisome Linker Gene Families in Mesophilic Red Algae’. *Nature Communications* 10(1):4823. doi: 10.1038/s41467-019-12779-1.

Lee, JunMo, Eun Chan Yang, Louis Graf, Ji Hyun Yang, Huan Qiu, Udi Zelzion, Cheong Xin Chan, Timothy G. Stephens, Andreas P. M. Weber, Ga Hun Boo, Sung Min Boo, Kyeong Mi Kim, Younhee Shin, Myunghee Jung, Seung Jae Lee, Hyung-Soon Yim, Jung-Hyun Lee, Debashish Bhattacharya, and Hwan Su Yoon. 2018. ‘Analysis of the Draft Genome of the Red Seaweed Gracilariopsis Chorda Provides Insights into Genome Size Evolution in Rhodophyta’. *Molecular Biology and Evolution* 35(8):1869–86. doi: 10.1093/molbev/msy081.

Nelson, David R., Khaled M. Hazzouri, Kyle J. Lauersen, Ashish Jaiswal, Amphun Chaiboonchoe, Alexandra Mystikou, Weiqi Fu, Sarah Daakour, Bushra Dohai, Amnah Alzahmi, David Nobles, Mark Hurd, Julie Sexton, Michael J. Preston, Joan Blanchette, Michael W. Lomas, Khaled M. A. Amiri, and Kourosh Salehi-Ashtiani. 2021. ‘Large-Scale Genome Sequencing Reveals the Driving Forces of Viruses in Microalgal Evolution’. *Cell Host & Microbe* 29(2):250-266.e8. doi: 10.1016/j.chom.2020.12.005.

Oliver, Aaron, Sheila Podell, Agnieszka Pinowska, Jesse C. Traller, Sarah R. Smith, Ryan McClure, Alex Beliaev, Pavlo Bohutskyi, Eric A. Hill, Ariel Rabines, Hong Zheng, Lisa Zeigler Allen, Alan Kuo, Igor V. Grigoriev, Andrew E. Allen, David Hazlebeck, and Eric E. Allen. 2021. ‘Diploid Genomic Architecture of Nitzschia Inconspicua, an Elite Biomass Production Diatom’. *Scientific Reports* 11(1):15592. doi: 10.1038/s41598-021-95106-3.

Osuna-Cruz, Cristina Maria, Gust Bilcke, Emmelien Vancaester, Sam De Decker, Atle M. Bones, Per Winge, Nicole Poulsen, Petra Bulankova, Bram Verhelst, Sien Audoor, Darja Belisova, Aikaterini Pargana, Monia Russo, Frederike Stock, Emilio Cirri, Tore Brembu, Georg Pohnert, Gwenael Piganeau, Maria Immacolata Ferrante, Thomas Mock, Lieven Sterck, Koen Sabbe, Lieven De Veylder, Wim Vyverman, and Klaas Vandepoele. 2020. ‘The Seminavis Robusta Genome Provides Insights into the Evolutionary Adaptations of Benthic Diatoms’. *Nature Communications* 11(1):3320. doi: 10.1038/s41467-020-17191-8.

Paajanen, Pirita, Jan Strauss, Cock van Oosterhout, Mark McMullan, Matthew D. Clark, and Thomas Mock. 2017. ‘Building a Locally Diploid Genome and Transcriptome of the Diatom Fragilariopsis Cylindrus’. *Scientific Data* 4:170149. doi: 10.1038/sdata.2017.149.

Sorokina, Maria, Emanuel Barth, Mahnoor Zulfiqar, Michiel Kwantes, Georg Pohnert, and Christoph Steinbeck. 2022. ‘Draft Genome Assembly and Sequencing Dataset of the Marine Diatom Skeletonema Cf. Costatum RCC75’. *Data in Brief* 41:107931. doi: 10.1016/j.dib.2022.107931.

Suzuki, Shigekatsu, Shuhei Ota, Takahiro Yamagishi, Akihiro Tuji, Haruyo Yamaguchi, and Masanobu Kawachi. 2022. ‘Rapid Transcriptomic and Physiological Changes in the Freshwater Pennate Diatom Mayamaea Pseudoterrestris in Response to Copper Exposure’. *DNA Research: An International Journal for Rapid Publication of Reports on Genes and Genomes* 29(6):dsac037. doi: 10.1093/dnares/dsac037.

Yamamoto, Kayoko, Takashi Hamaji, Hiroko Kawai-Toyooka, Ryo Matsuzaki, Fumio Takahashi, Yoshiki Nishimura, Masanobu Kawachi, Hideki Noguchi, Yohei Minakuchi, James G. Umen, Atsushi Toyoda, and Hisayoshi Nozaki. 2021. ‘Three Genomes in the Algal Genus Volvox Reveal the Fate of a Haploid Sex-Determining Region after a Transition to Homothallism’. *Proceedings of the National Academy of Sciences of the United States of America* 118(21):e2100712118. doi: 10.1073/pnas.2100712118.

Zepernick, Brittany N., David J. Niknejad, Gwendolyn F. Stark, Alexander R. Truchon, Robbie M. Martin, Karen L. Rossignol, Hans W. Paerl, and Steven W. Wilhelm. 2022. ‘Morphological, Physiological, and Transcriptional Responses of the Freshwater Diatom Fragilaria Crotonensis to Elevated pH Conditions’. *Frontiers in Microbiology* 13:1044464. doi: 10.3389/fmicb.2022.1044464.

Zhang, Zehao, Junhao Wang, Xiaoqian Zhang, Xiaowei Guan, Tian Gao, Yunxiang Mao, Ansgar Poetsch, and Dongmei Wang. 2023. ‘ChIP-Based Nuclear DNA Isolation for Genome Sequencing in Pyropia to Remove Cytosol and Bacterial DNA Contamination’. *Plants (Basel, Switzerland)* 12(9):1883. doi: 10.3390/plants12091883.
